# Guiding Discovery of Protein Sequence-Structure-Function Modeling

**DOI:** 10.1101/2023.07.14.548822

**Authors:** Azam Hussain, Charles L. Brooks

## Abstract

Protein engineering techniques are key in designing novel catalysts for a wide range of reactions. Although approaches vary in their exploration of the sequence-structure-function paradigm, they are often hampered by the labor-intensive steps of protein expression and screening. In this work, we describe the development and testing of a high throughput *in silico* sequence-structure-function pipeline using AlphaFold2 and Fast Fourier Transform docking that is benchmarked with enantioselectivity and reactivity predictions for an ancestral sequence library of fungal flavin-dependent monooxygenases. The predicted enantioselectivities and reactivities correlate well with previously described screens of an experimentally available subset of these proteins and capture known changes in enantioselectivity across the phylogenetic tree representing ancestorial proteins from this family. With this pipeline established as our functional screen, we apply ensemble decision tree models and explainable AI techniques to build sequence function models and extract critical residues within the binding site and the second sphere residues around this site. We demonstrate that the top-identified key residues in the control of enantioselectivity and reactivity correspond to experimentally verified residues. The *in silico* sequence-to-function pipeline serves as an accelerated framework to inform protein engineering efforts.

## Introduction

Protein engineering is the design of proteins for improved or unique fitness, where fitness can describe any property of the protein including reactivity, enantioselectivity, or thermostability^1,2^. Approaches to protein engineering explore various aspects of the sequence-structure-function paradigm. One of the most popular and successful strategies is the use of directed evolution, where libraries of variants are constructed by mutating the wildtype sequence^1,3^. Variants with high fitness are selected and used in further iterations to create new highly fit enzymes. Directed evolution takes advantage of the vast exploration of the sequence landscape with sequencing and assaying at scale. However, directed evolution is labor intensive and does not leverage 3D structural information that may guide sequence design and prediction of protein function. Rational design, on the other hand, takes advantage of the protein structure and interactions between protein and its native substrate to engineer the protein^4,5^. However, it requires an accurate model of not only the protein structure but also interactions and mechanistic insight into substrates of interest. A high-throughput computational protocol is required that can offer rapid and informed discovery of new variants to explore while bridging the gap between sequence, structure, and function.

AlphaFold2^6^ (AF2), the top-performing structure prediction method in CASP14^7^, has made it possible to rapidly generate high-quality structures of novel sequences. Enormous structure libraries such as the AF2 database of protein structure predictions^8^ and ESM Metagenomic Atlas^9^ demonstrate the possibility to utilize and explore novel prediction methods at scale^10^. Previously, state-of-the-art structural prediction methods were limited to the domain of hours to days^11^, but examples such as ColabFold^12^, DMPFold2^13^, ESMFold^9^, and RoseTTAfold^14^ have made state-of-the-art structure prediction accessible in minutes to hours. ColabFold uses a modified multiple sequence alignment (MSA) pipeline utilizing MMseqs2^15^, demonstrating that AF2 can be repurposed for speed and accessibility. We describe an approach that also utilizes a modified MSA generation pipeline and enables us to quickly explore a target enzyme family.

Multiple approaches exist for associating structures generated in such a manner with their function^16^. High throughput methods to relate protein and ligand structure to affinity or reactivity generally use empirical models and quantitative structure-activity relationships^17^. However, predictions that provide geometric and functional insight rely on protein-ligand docking and the prediction of binding affinity^18^. Molecular docking has made it possible to predict with reasonable accuracy the binding poses and affinities of substrates in protein structures, creating the opportunity to predict protein function *a priori* from the binding pose and docking score of the ligand. To dock an array of multiple ligands to hundreds of protein structures, a high throughput docking approach is needed with an accurate scoring method.

In the work presented here, we describe a high throughput protocol for building structure-function models *in silico* that leverages AF2^6^ and GPU-accelerated Fast Fourier Transform-based docking (FFTDock)^19^ as implemented in CHARMM^20^ utilizing the physical forcefields from the CHARMM36^20,21^ and CGenFF^22^ forcefield efforts. To assess the validity of the high throughput structure and docking pipeline, we focus on a model of catalysis by fungal flavin-dependent monooxygenases (FDMOs). TropB is an FDMO that carries out oxidative dearomatization, a useful reaction in organic synthesis that exhibits high site and stereoselectivity across a variety of resorcinol substrates^23^. AfoD, AzaH, and SorbC are related FDMOs that also possess unique reactivities and site-selectivities with relatively minor changes in the steric and electronic environments of their substrates^23,24^. Structural models and molecular modeling have been utilized to probe function, mechanism, and rational engineering^24–27^ in this family of FDMOs as well. For example, a mechanistic study of TropB revealed that the face of the ligand presented towards the activated FAD cofactor leads to hydroxyl group addition on that face, suggesting the use of molecular docking to elucidate stereochemistry and reactivity. Additionally, ancestral sequence reconstruction (ASR) of mammalian FDMOs has been used to find stable ancestors and learn important structural features^28^. More recent work has demonstrated the efficacy of using ancestral sequence resurrects to determine key residues controlling stereoselectivity in fungal FDMOs TropB, AfoD, and AzaH^29^. In the work we describe below, we demonstrate the protocol we lay out in the following can infer *a priori* protein enantioselectivity and reactivity from interactions between protein and ligand structural models and predict the effect of known key stereochemical switches^29^.

Our approach builds upon previous exploration of structure-function models to guide design. A previous pipeline^30^ for modeling ASR enzymes illustrated the use of MODELLER^31^ to apply high-throughput homology structure modeling to a family of double-stranded RNA binding enzymes. MODELLER requires high CPU usage to scale to larger phylogenies and the protocol relied on pre-trained models to assign binding affinity to the structures. Wong et al.^32^ demonstrated the use of AF2 database structures^8^ and AutoDock Vina^33^ to screen anti-bacterial compounds against *E. coli* essential proteins and demonstrated the use of machine learning (ML) rescoring functions to slightly improve binding affinity predictions. AutoDock Vina is more computationally expensive than FFTDock and alone was unable to predict top binders. Wijima et al.^34^ demonstrated *in silico* enzyme design of enantioselective enzymes by using multiple independent molecular dynamics simulations in a high-throughput fashion (HTMI-MD)^35,36^, but this approach relies on a single starting crystal structure, docking of predefined R and S orientations and brute force molecular dynamics simulations, and hence is of limited scalability. AlphaFill^37^ is an algorithm that matches structures with cofactors and ligands in the PDB library to models in the AF2 structure database and uses the YASARA^38^ forcefield to minimize transplanted small molecules into AF2 models. We describe a similar approach of transplanting the FAD cofactor into our predicted structures using the CHARMM36^21^ and CGenFF^22^ forcefields. Nevertheless, these efforts serve as inspiration for the work we describe below.

Ultimately the driving goal in structurally characterizing enzymes is to elucidate the key determinants of function. However, given the large number of predictions generated for multiple ligands and a large sequence-structure space, it can be difficult to infer the residues to target in the design of a better biocatalyst. ML approaches have been applied in the context of directed evolution to map enormous sequence-fitness landscapes, speeding up directed evolution with a more informed selection of mutations^39,40^. In particular, the ML framework of gradient-boosted trees has been previously used to fit enantioselectivity to enzyme properties^41^. We describe a generalizable approach to fit a sequence-function model using an ensemble of decision tree methods, by representing the sequence data in the tabular form of an MSA. We then identify residues that control reactivity and stereochemistry using SHapley Additive exPlanations (SHAP)^42^. SHAP is an approach to linearly approximate the features determining a model’s prediction and has been widely adopted in the field of explainable artificial intelligence (AI)^43^. SHAP has been previously used to understand the role of features such as composition, property, and nucleotide type in mRNA modification site prediction^44,45^, and to highlight key functional groups in small molecule potency predictors^45^. We demonstrate its application to protein sequence-function analysis, with SHAP values assigning each amino acid to a stereochemistry and reactivity contribution. SHAP analysis of key residues from *in silico* sequence-function pairs will serve as a rapid and reasonably accurate step to guide protein engineering efforts.

## Results and Discussion

Our objective in the current work is to describe and demonstrate a high-throughput framework for exploring and optimizing biocatalytic enzyme function through the combination of modern structure prediction, ligand docking and refinement, and ML-based sequence-function modeling. This framework and workflow for sequence-structure-function prediction is illustrated in Figure 1, where we show the components of our predictive scheme comprising two basic elements: structure-based function prediction (reactivity and stereoselectivity) followed by sequence-based ML to identify key protein residues responsible for determining reactivity and stereoselectivity. These two components constitute a method to guide rational design of novel biocatalysts, to rationalize observed sequence-function screens, and to direct and inform rational design approaches based on directed evolution.

**Figure 1.**
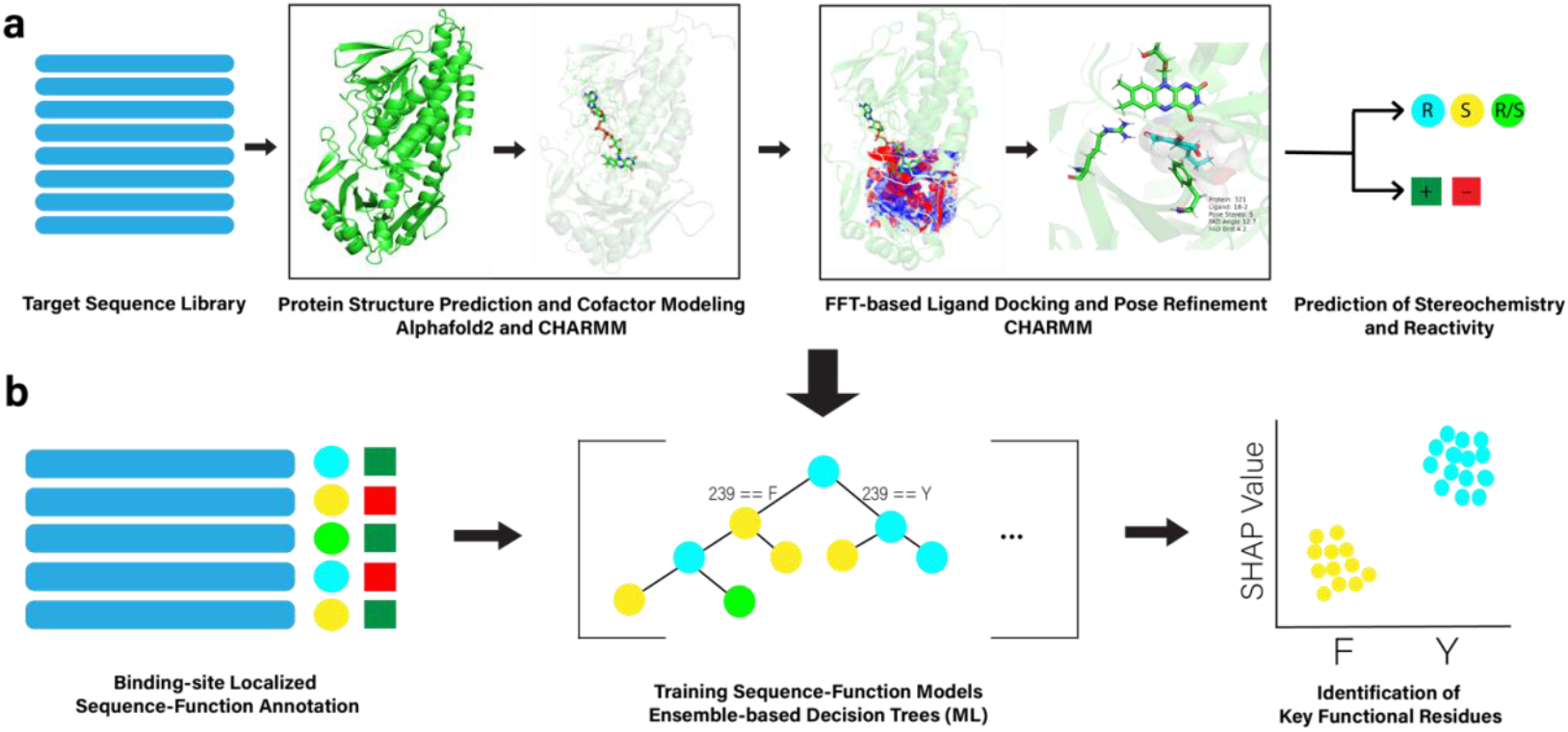
Protein sequence-structure-function pipeline. a) Protein structures are predicted from a target sequence library using a modified AF2 protocol. Following, any cofactors are modeled into structures (in the present case FAD) and ligands are docked into protein structures using FFT-based docking^19^. The resulting structures are utilized to predict the expected stereochemistry and reactivity of each member of the target sequence library. b) Residues representing aligned sequences from the first and second shell around the binding site are arranged into an MSA and combined with the sequence-function predictions from above as input into an ML model. Using an ensemble of random forests and gradient-boosted tree methods, results for the enantioselectivity and reactivity are learned. SHAP analysis is then employed on these learned models to identify key residues to guide engineering.

In the following we will illustrate this methodology through application to the rational exploration of an ancestral sequence reconstruction (ASR) of a family of fungal FDMOs in which we: i) create structural models of extant sequences, predicted ancestor sequences, and the first-alternative sequences from the ASR, including the incorporation of cofactor FAD into all structures using a modified AF2 prediction methodology^6^ and molecular modeling, ii) dock a panel of four ligands, which are part of the experimental screen of these proteins for catalytic activity and stereochemistry outcome^29^, using FFT-based ligand-receptor docking methods^19^, iii) create classifiers based on these structural models and docking approach to predict both reactivity and stereochemistry. We then use this framework in conjunction with sequence-based ML methods to identify key binding/active site and second-sphere residues responsible for the control of stereochemical switching of products. The elements i-iii are tested and evaluated by comparison of predictions for known structures (although unknown to the AF2 prediction framework), assessment of docking poses compared with known ligand positioning within the family of extant proteins, and through comparison of predictions for a set of 67 enzymes previously screened experimentally for reactivity and stereochemistry of products. The final component of our workflow is evaluated by comparison of identified key residues with experimental mutational studies. In what follows we discuss the elements described above and present our findings in both assessing the methods and in their application to annotating the reactivity and stereochemistry of the predicted ASR.

### Fast High-Fidelity Models with Alphafold2 Using Consensus Sequence Hits

The AF2^6^ models built using MSAs from the consensus sequence hits (see Methods below) showed good agreement with TropB and AfoD crystal structures (Figure 2). After alignment with TM-align^46^, the TropB AF2 model and crystal structure had a Cα root mean square deviation (RMSD) of 0.91 Å (Figure 2a) and AfoD had a Cα RMSD of 0.99 Å (Figure 2c). For consistency we use the residue numbering and amino acid lettering of TropB to refer to residues across enzymes (see Supplementary Data 1 for list of extant and ASR enzymes). Key binding site residues R206 and Y239 are identically positioned with the crystal structure in TropB (Figure 2c) and AfoD (Figure 2d). R206 and Y239 do not significantly change position compared to the QM/MM refined model^27^, suggesting that docking to the apoprotein can recapitulate the correct pose and stereochemistry. While we did not limit the template space searched, we observed that even with dummy templates a high agreement was obtained for the crystal structures of AfoD and TropB, with an overall Cα RMSD of 1.528 Å for TropB (Table S1) and 1.525 Å for AfoD. This suggests that our modifications of the AF2 prediction pipeline with limited sequences and templates are robust. Predictions for AfoD and TropB were on par with other protein structure prediction web servers, despite our smaller preconditioned library for MSA generation. This suggests that the initial sequences and templates found from constructing the ancestral tree or hits from the consensus sequence can be repurposed for rapid structure prediction aided by GPU-based model inference and AMBER^47^ relaxation.

**Figure 2.**
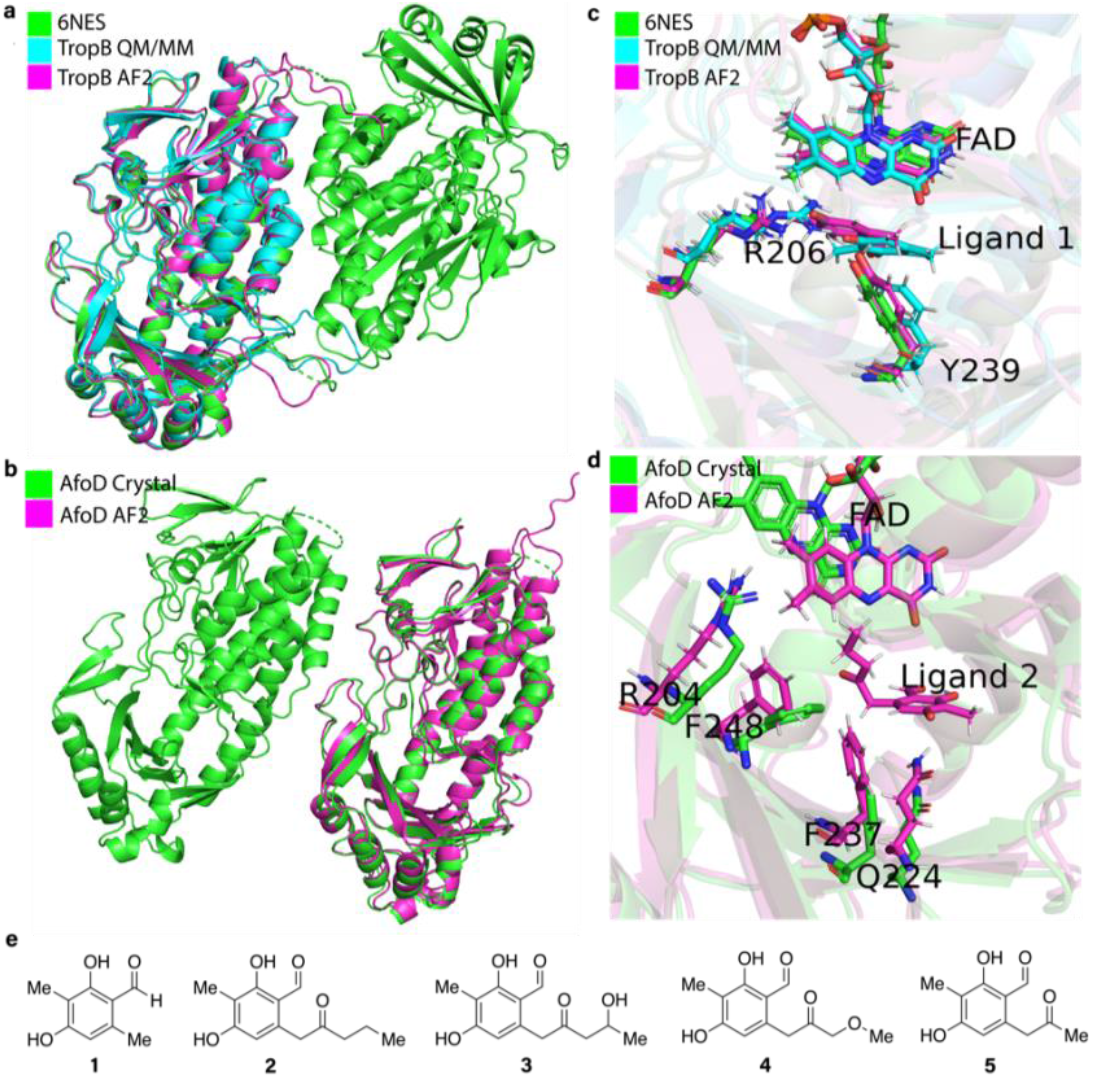
TropB and AfoD AF2 Comparison with Crystal Structures. a) TropB crystal structure (green, RCSB PDB ID: 6NES) structurally aligned to AF2^6^ model (purple) generated using consensus hits. b) AfoD crystal structure (green, RCSB PDB ID: 7LO1) structurally aligned to AF2 model (purple). c) TropB crystal structure (green), AF2 model (purple), and a QM/MM refined structure (cyan) docked with native substrate **1** using Flexible CDOCKER^27^. AF2 model docked with **1** using FFTDock and protein minimization (residues Y239 and R206 are indicated). Docked pose in AF2. d) AfoD crystal structure and AF2 structure docked with FAD and **2** using FFTDock and protein minimization. e) TropB native substrate **1**, model substrate **2**, AzaH native substrate **3**. Substrates **2-5** were previously tested against the ancestral FDMO library for stereoselectivity^29^.

### Predicted Structures Show High Predicted Accuracy with Consensus MSA Library

Because of the increased speed of prediction resulting from the consensus library method we implemented, obtaining structural models for thousands of ancestral and extant proteins is possible with modeling on the order of minutes for each structure. To judge the quality of these structures we utilize the per residue predicted Local Distance Difference Test (pLDDT)^48^ score from AF2. The pLDDT score represents a measure of model confidence for the prediction. The average pLDDT score across the 830 structural models predicted for the ASR sequence library was 93.2, indicating confidence in high-quality structures (Figure S2). It is worth noting that in some of the most distant ancestors, for example ancestor 278 at the root of tree, very few binding site residues are in common, and an overall pLDDT of 64.1 suggests a structural model of somewhat lower quality is predicted in this case (Figure S3). With the original AF2 MSA search pipeline, we observed an overall pLDDT of 69.3 for ancestor 278 (Figure S4), indicating that the consensus library is not a key contributor to the lower confidence. The consensus structural model for ancestor 278 still maintains higher accuracy in the more conserved central core (Figure S3), suggesting that even lower-scoring models are useful in providing mechanistic insight.

### Positioning Cofactor FAD in its Binding Pocket

In the family of fungal FDMOs the cofactor FAD is known to exist in both an IN and an OUT conformation^27^, related to the mechanistic role of FAD in both catalyzing the chemistry on substrate molecules in the IN conformation and recycling its oxidative state by interacting with external cofactor NADPH (or NADH) in the OUT conformation. Consequently, the FAD cofactor was positioned in the binding pocket for the IN conformation (see Methods for details) because it is most like the catalytically active conformation with C4α-hydroperoxyflavin^25^. Docking ligands based on the IN conformation should capture near-native poses that are poised to adopt the correct stereochemistry in their products. After FAD incorporation, the interaction energy between the FAD cofactor and the AF2 structure of TropB was -198.90 kcal/mol suggesting a highly favorable interaction and correct modeling of the FAD pocket. The majority of the sequence library had the FAD cofactor pocket successfully modeled and incorporated into the structures, based on our observations of similar interaction energies between the proteins and the FAD cofactor. An average of -210.8 kcal/mol across all structures was obtained (Figure S5). Hydrogen bonding interactions dominate between FAD and the protein with an average electrostatic interaction energy of -160.4 kcal/mol and an average van der Waals interaction energy of -50.4 kcal/mol. Thus, incorporation of the cofactor into our predicted models was relatively straightforward and gave us confidence in docking ligands into the structures predicted from our AF2 pipeline.

### Fast Rigid Receptor Docking and Stereochemistry Prediction

The ligands (ligands **2-5** illustrated in Figure 2e) were docked into our predicted cofactor-integrated AF2 protein structures utilizing the GPU-accelerated FFTDock protocol^19^ available in CHARMM^19^. In a few seconds thousands of poses can be generated and scored. Because of the use of a soft grid (see Methods), many of these poses had some overlap with protein backbone atoms and needed further refinement. We explored various strategies for rescoring these poses, by minimizing them further in the FFTDock grid, minimization in an explicit protein representation, and using the starting poses as conformers for simulated annealing based flexible ligand docking^19^ Simulated annealing allows for further exploration of ligand conformational space, while explicit protein gives the most accurate but computationally expensive representation of the protein.

To benchmark different rescoring strategies, we compared predicted stereochemistry from docking against known stereochemistries of **2-5** in the 67 enzymes of the ancestral FDMO library^29^ (see also Figure 2e), using the angle of the clustered poses relative to the FAD cofactor (see Methods, Figure S6). We calculated the overall predicted stereochemistry for an enzyme-ligand pair by treating the system as a Boltzmann-weighted ensemble of R and S mesostates (see Eqn 1 in Methods below). This strategy allows multiple top hits to contribute to the final stereochemistry prediction (i.e., a consensus), reducing the influence of highly ranked outlier clusters. Simulated annealing and minimization in a fixed protein environment achieved similar stereochemical accuracies ranging from 55-70% across all experimentally active protein-ligand pairs, similar in performance to traditional docking methods in recapitulating native poses^49^ (Figure S7). A general trend we observed was that using a lower dielectric constant improved prediction accuracy and Matthew’s correlation coefficient (MCC)^50,51^ as is illustrated in Figure S8. The top-performing docking approach was minimization of FFTDock poses in explicit protein with an R-dependent dielectric constant of 0.75, giving an accuracy for stereochemistry prediction across all protein-ligand pairs of 73% and an MCC of 0.51 (Table S2). The MCC score indicates the successful prediction of both R and S poses. Finding the consensus stereochemistry by averaging across ligands **2-5** leads to slightly increased robustness with this selected rescoring strategy yielding an overall accuracy of 77% and an MCC of 0.65 (Figure S9).

As is evident in Figure 3, from the optimal scoring/stereochemistry prediction protocol just discussed we find very good agreement between the predicted and observed screening results. Moreover, where reliable docked poses are found for proteins that appear as unreactive in the experimental screen, which may be indicative of substrates that fail to convert but may also result from sub-optimal conditions used in screening for a given substrate-protein pair, we are able to predict the anticipated stereochemistry of the product. While this may be a useful outcome, because we are aiming to predict the stereochemistry and reactivity of a large family of proteins to inform choices regarding which sequences within the ASR to resurrect and test through protein expression and screening, we developed a machine learning model to enable us to predict substrate-protein pair reactivity in the specific screen being employed.

**Figure 3.**
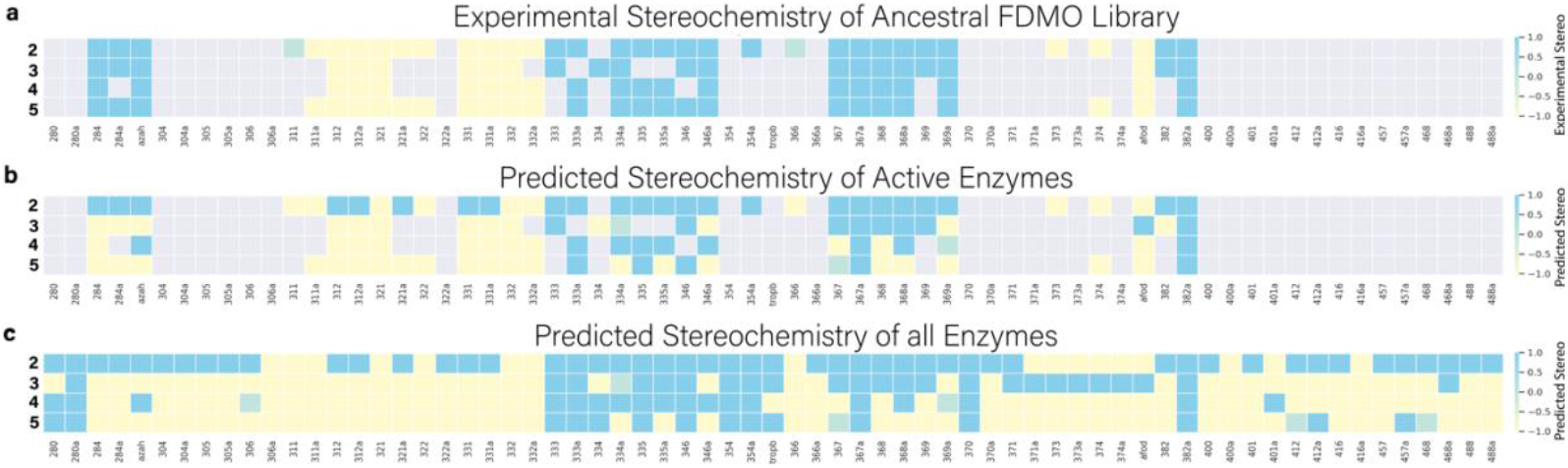
Stereochemistry Predictions. a) Experimentally observed stereochemistry for the ancestral FDMO library from previous work^29^. The stereochemistry of ligands **2-5** is depicted in rows along the y-axis and the screened proteins in the library are the columns (x-axis). Grey squares indicate proteins for which low/no reactivity is observed in the screen, yellow squares indicate S stereochemistry, blue squares indicate R, and green squares indicate promiscuity of enantiomer with both R and S (R/S) stereochemistry being observed. b) Predicted stereochemistry from the sequence-function screen for the reactive subset of the ancestral FDMO library. c) Predicted stereochemistry from sequence-function model for all enzymes in ancestral FDMO library.

### Prediction of Reactivity with Docking Energy and ML Rescoring

Docking metrics representing possible features correlating with enzyme reactivity were extracted from the top-ranked ligand pose. FAD distance and FAD angle were measures of the ligand proximity and orientation necessary for reacting with the c4α-hydroperoxyflavin^25^. Anion distance was chosen to represent the importance of R206 and Y239 in positioning the substrate in TropB^27^. Docking energy efficiency and Pafnucy^52^ predicted pK_d_ efficiency were chosen as measures of the overall binding affinity of the protein and ligand, utilizing information from the CHARMM36^21^ and CGenFF^22^ forcefields and a convolutional neural network (CNN) trained on binding affinities from PDBBind^53^. FAD distance was determined to be non-predictive of reactivity, with a p-value greater than .10 in 3 out of 4 ligand-specific logistic regressor models. The ligand-specific logistic regression models achieved an accuracy of 75.4% and an MCC of 0.50 across all ancestral FDMO protein-ligand pairs (Table S4). We explored averaging metrics across ligands to create a consensus predictor of reactivity. The consensus logistic model obtained an accuracy of 79.1% and a MCC of .57, based on predicting conversion in a protein that displayed any reactivity with ligands **2-5**. The predicted pK_d_ from Pafnucy and docking energy were identified as the top predictors of reactivity (Table S3), demonstrating the utility of ML-based scoring functions as rescoring methods in combination with docking scores.

We have established the fidelity and accuracy of each of the components of our pipeline in the above discussion, showing that very good predictions can rapidly be obtained for the structure of the protein and the pose of the ligand, and that from these the prediction of stereochemistry and reactivity can be achieved. Thus, combining the components of our sequence-structure-function pipeline we are now in a strong position to “annotate” sequences of unknown structure and function and serve to inform experimental studies seeking to discover novel protein sequences as a basis for biocatalyst design (or redesign). We also note that in principle, while not directly explored in this study, this pipeline model can be applied to the “functional annotation” of any collection of protein sequences for which high-throughput predictions of protein-bound ligand structures can be utilized to establish metrics from which reactivity and function can be inferred. We move on to explore the annotation of the constructed ancestral phylogeny of fungal FDMOs related to TropB, AfoD, and AzaH^29^ and to identify those residues within the active/binding site and in the second sphere of residues around this set that are key determinants to the functional outcome for each sequence.

### Annotation of All Sequences

Average stereochemistry and consensus reactivity predictions for ligands **2-5** were generated for the full ASR sequence library and are shown in Figure 4 and available in Supplementary Data 1. Our predictions suggest that clades are grouped by similar stereochemistry and reactivities, indicating the structure-function pipeline can capture the connection between sequence and functional similarity. The TropB, AfoD, and AzaH clades were classified as reactive, and the reactivity predictor identified other potential clades to be reactive for substrates **2-5**, which provides starting points for further exploration of the phylogenetic tree. The ancestral FDMO library was used to experimentally demonstrate a stereochemistry shift from S to R in the TropB (R) clade, and an R to S transition in the AfoD (S) clade. The shift in stereochemistry is controlled by the identity of residue 239, with F239 promoting S stereochemistry, and Y239 promoting R stereochemistry^29^. The TropB clade is predicted to contain two subfamilies of R and S enzymes consistent with the residue 239 F/Y switch. An intermediate clade between TropB and AzaH, containing ancestor 455, exhibits R stereochemistry and reactivity while possessing phenylalanine at residue 239 like AzaH. This demonstrates how the pipeline can be used to identify novel sequences to guide exploration, where future studies could focus on ancestors near 455 as a bridge between the unique stereo-control mechanism of AzaH^29^ and the residue 239 F/Y switch in AfoD and TropB. We discuss the previously explored stereo-control mechanisms of TropB, AfoD, and AzaH^29^ in further detail and the capacity of the proposed pipeline to capture these mechanisms.

**Figure 4.**
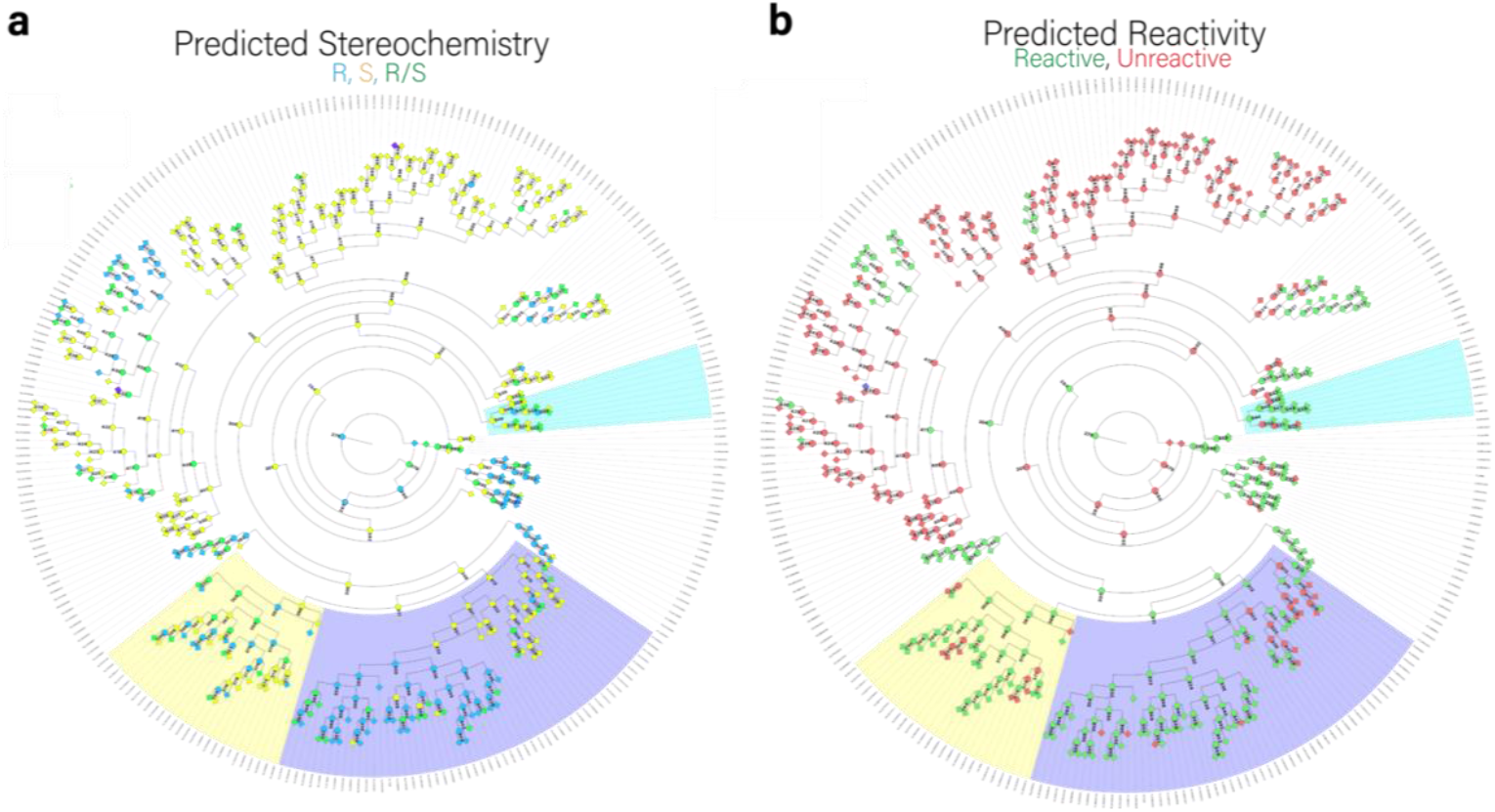
Phylogenetic tree with predicted stereochemistry and reactivity from the sequence-structure-function protocol. a) Phylogenetic tree with mean predicted stereochemistry results using FigTree. Ancestors and extant sequences are colored in by the average stereochemistry of ligands **2-5**, with S as yellow, R as blue, and R/S as green. The TropB (R) clade is highlighted in blue, the AfoD (S) clade is highlighted in yellow, and the AzaH (R) clade is highlighted in cyan. b) Phylogenetic tree with predicted reactivity with averaged metrics. Ancestors and extant sequences are colored by the predicted reactivity for ligands **2-5**, with green indicating reactive and red indicating unreactive.

### Predictions in the AfoD, TropB, and AzaH clades

The observed binding pose in the TropB clade across all substrates is similar to previous work characterizing TropB’s native substrate utilizing varying docking approaches^25,27^. The R-binding pose in ancestors 333, 334, 335, and 346 with Y239 is consistent with the previously observed binding pose of TropB (Figure S11). The S-binding pose in ancestors with F239 is found to involve a rotation of the resorcinol core around the anion axis, maintaining the anion hydrogen bonding interaction with R206 (Figure S12). For the AfoD clade, in which we observe the stereochemistry switch from R to S (Figure S13), ancestor 369 has an R binding pose consistent across all the R-producing enzymes. For AfoD and its S-producing ancestors with the F239 mutation, a V250F substitution can be found that blocks Arg206 from accessing the substrate and creates an active site surrounded by three phenylalanine residues (Fig 2d). Thus, the binding pose is entirely flipped but still leads to the S-product. This unique orientation of ligands in AfoD and its S-producing ancestors informs the exploration of AfoD-specific design targets.

AzaH performs R stereochemistry but possesses phenylalanine at residue 239, leading to a separate stereo-control mechanism. The AzaH clade is predicted to be predominately S, and AzaH on average is predicted to be racemic, with R stereochemistry for 2/4 ligands (Figure S14). The top ligand clusters of AzaH for **3** are dominated by low scoring S oriented representative poses, most likely favored by F239. Simulated annealing and protein minimization at various epsilon (the dielectric constant used in the refinement and scoring of ligand poses) still lead to identification of ligands binding to AzaH as S or racemic. This suggests that the more complex stereo-control mechanism of AzaH is not fully captured by the modeling protocol and may need to be explored further using flexible docking or docking to an ensemble of AF2 structures to better delineate the conformational space of the protein receptor. From the results for AzaH, one may argue that the protocol simply captures the role of residue 239 alone. However, we demonstrate with mutations to residue 239 and sequence function modeling the capacity of the model to explore a more complex stereo-control mechanism through exploration of the rest of the sequence library.

### Prediction of Residue 239 Mutations

Residue 239 plays a key role in stereochemistry determination, where F239 promotes S stereochemistry and Y239 promotes R stereochemistry. F239Y in AfoD promotes the catalysis of a racemic product distribution^29^ and the functional screen should capture this effect. We tested the functional screen in predicting the effect of mutation to this key residue by running the full screen on F239Y and Y239F variants of the whole ASR library. In the ancestral FDMO library (the 67 enzymes assayed with **2-5**), we observed 65.6% of protein-ligand pairs change average stereochemistry from the wildtype stereochemistry to racemic or the opposite stereochemistry (Figure S15 a-b). 61.4% of proteins in the ancestral FDMO library with the F239Y mutation changed stereochemistry and 73.9% with the Y239F mutation. This indicates that the functional screen captures the significant role residue 239 plays in stereochemistry, and the observation that mutation of residue 239 alone is insufficient to fully change stereochemistry. In the entire sequence library, sequences with mutation Y239F resulted in a 22.7% decrease in the number of predicted R class, and sequences with mutation F239Y had a 50.3% decrease in the number of predicted S class (Figure S15 c-d). This suggests that mutation to residue 239, while key in a majority of the tree, is not able by itself to fully control stereochemistry. We then applied ML models to identify potential residues besides 239 that contribute to the stereo-control mechanism.

### Sequence-Function Modeling with Random Forest and Gradient Boosted Trees

We trained multiple sequence-function models on all sequence-function pairs to predict average stereochemistry and consensus reactivity. An MSA of binding site and previously unexplored second shell residues of the 830 sequences in the full ASR library was used as an input to the model (Figure S16), in order to predict the previously obtained average stereochemistry and consensus reactivity with ligands **2-5**. To determine the best ML architecture for this problem we used the mljar AutoML framework^54^ to train models ranging in complexity from linear models, ensemble-based decision tree methods, support vector machines, and feedforward neural networks. Ensemble tree-based methods were most successful in predicting stereochemistry and reactivity and demonstrated that the sequence-structure-function pipeline predicted properties that could be mapped to the original sequence (Figures S17-S18). We then used mljar in perform mode to create multiple hyperparameter-tuned models for the CatBoost^55^, XGBoost^56^, and Random Forest^57^ algorithms which slightly increased performance relative to the default models (Figures S19-S20). The best-performing model for stereochemistry prediction was a hyperparameter-tuned CatBoost model that yielded an accuracy of 74.3% and a macro average F1 score of .547. The best-performing model for reactivity prediction was a hyperparameter-tuned XGBoost model with an accuracy of 87.7% and an F1 score of .88.

### Key Features from SHAP Analysis

The hyperparameter-tuned models did not have significantly different performances but varied in agreement among the top residues by mean absolute SHAP value (Figures S21 and S23). To overcome selection bias from choosing a top model, we used the consensus among all folds of all models by averaging the normalized mean absolute SHAP values to suggest consensus top residues (Figures S22 and S24). We observed that for both stereochemistry and reactivity prediction, the first highest-ranked residues were the binding pocket, but multiple unexplored second-shell residues in a specific region played a key role in the prediction (Figure 5 a,c). Residue 239 was scored as the highest contributor to stereochemistry prediction, suggesting that it plays a significant role not only in the AfoD and TropB clades but across the entirety of the sequence library. The SHAP dependence plot for residue 239 (Figure 5b) allows for easy interpretation of the F/Y switch, and dependence plots can be used for other top features to guide design strategies. Interestingly, M54, a previously unexplored residue, was scored as the highest contributor to reactivity prediction across all sequences, with residue 239 scored as the second most important feature. The side chain of residue 54 is located near the FAD cofactor, and changes at position 54 would indirectly affect ligand binding through the interaction with FAD. According to the dependence plot in Figure 5d, V54 significantly negatively impacts reactivity and suggests an I/V reactivity switch as the library has 48.6% V and 35.9% I. V54 is associated with a lower average docking score and Pafnucy pK_d_ (Figure S25 a-b). This suggests a novel approach to engineering selectivity by engineering residues around the FAD cofactor to modify the ligand pocket and would demonstrate an interesting avenue to explore.

**Figure 5.**
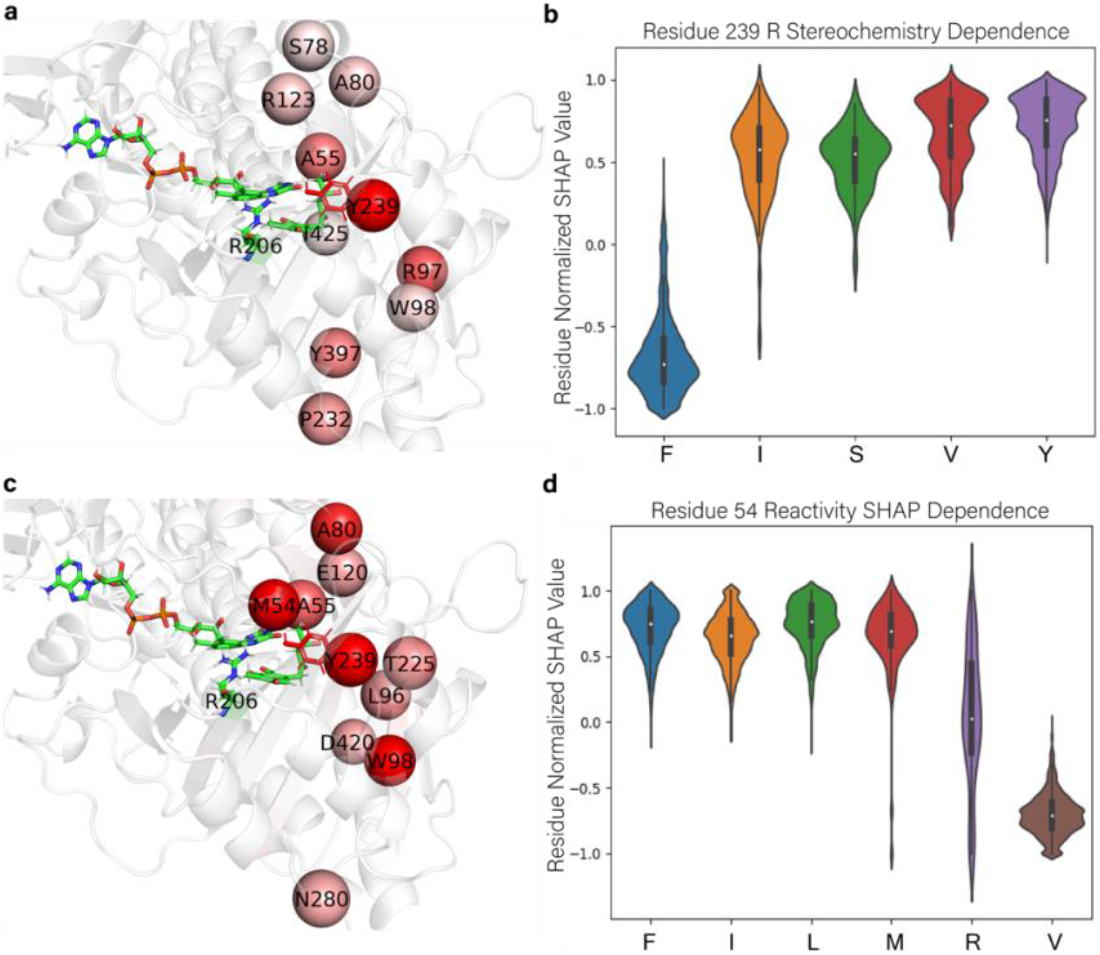
Key residues for stereoselectivity and reactivity control identified by SHAP analysis. a) Top 10 residues by mean absolute normalized SHAP value across all generated folds of all models for prediction of stereochemistry in TropB docked with **3**. The deep red spheres indicate a high mean absolute SHAP value, with Y239 having the largest value. b) SHAP dependence plot of residue 239 for the R class with normalized SHAP contribution of residue 239 across all folds of all models for full sequence library. c) Top 10 residues by mean absolute normalized SHAP value across all generated folds of all models for prediction of reactivity. M54 has the largest mean absolute SHAP value. d) SHAP dependence plot of residue 54 across all folds of all models for full sequence library for prediction of reactivity.

## Conclusion

Alphafold2 has revolutionized protein structure prediction, not only in terms of accuracy but also in speed and accessibility. Together with state-of-the-art docking approaches and machine learning frameworks, we are moving closer to the successful prediction and understanding the function of proteins *in silico* and, clearly, to a facile means of exploring key problems in the engineering of enzymes. In the work presented here, we have developed a framework for the establishment of a sequence-structure-function pipeline for the prediction of protein structures, function, and key residues. The approach is generalizable to many protein systems, and we successfully demonstrated for the case of fungal flavin-dependent monooxygenases that this framework and the specific pipeline we developed for this application can not only recapitulate enantioselectivity and reactivity with good accuracy, but also that it can guide new approaches to engineering. The protocol can annotate stereochemistry and reactivity of unexplored enzymes and provide more informed selection of novel enzymes to explore. In addition, through the sequence-function model we are able capture roles of more distant residues that would normally elude a first pass in which active/binding site adjacency only are considered. We anticipate that sequence-structure-function frameworks based on the ideas we present and discuss here will serve a significant role in informing future studies aimed at the engineering and design of new proteins for specified functional purposes.

## Acknowledgements

The authors acknowledge the efforts of Chang-Hwa (Chad) Chiang and Professor Alison Narayan for efforts to resurrect and screen protein sequences from the family of fungal flavin-dependent monooxygenases in their laboratory and to Dr Troy Wymore for his work in constructing the ASR for proteins in this family. The National Institutes of Health is acknowledged for financial support under grant GM130587.

## Methods

### Sequence Library

The wild-type sequence library consisted of 277 extant flavin-dependent monooxygenase sequences, 276 maximum likelihood ancestral resurrect sequences, and 276 alt-all ancestral resurrect sequences as previously described^29^. Of these, 67 were previously expressed and experimentally assayed for stereochemistry and conversion denoted as the ancestral FDMO library These sequences formed the basis for training and testing our pipeline and are included in Supplementary Data 1.

### Model generation with Alphafold2

A consensus sequence from the MSA of the 277 extant sequences used to perform ASR^29^ (Supplementary Data 2) was generated using HHconsensus from HHsuite3^58^, with match states in columns with less than 50% gaps. AF2 v2.0’s data pipeline, model weights, and inference script were used. AF2’s data pipeline was used to generate MSAs from the consensus sequence and the MSAs were combined into a FASTA formatted set of 84,572 sequences, representing the consensus sequence hits. This database was used to generate AF2 models of the ancestral sequences by replacing the standard data pipeline for the feature dictionary with a JackHMMER search on the consensus sequence hits. The standard AF2 MSA pipeline’s BFD database consists of over 2.5 billion sequences, with HHblits on the BFD database being CPU limited and highly I/O intensive. Using the consensus sequence hits reduces MSA generation from hours/days to under a minute. The MSA of the top 10000 hits was used with HHsearch on the PDB70 database^58^ to find templates with AF2’s template featurizer. The model generation step used monomer model 1 with 1 ensemble and default Amber relaxation constants.

### Minimization and Addition of FAD Cofactor

The AF2 models were superposed with TM-align^59^ to the previously generated QM/MM refined chain A of RCSB PDB 6NES docked with 3-methyl-orcinaldeyde (3MO)^27^. The superposed structures were represented with CHARMM in vacuum using the pyCHARMM^60^ package. The superposed structures were minimized using 1000 steps of steepest descents (SD). Next, the FAD cofactor present in the QM/MM structure was added to the models. The FAD cofactor was minimized in the structure using successive rounds of SD and Adopted Basis Newton Raphson (ABNR) minimization, with more and more of the protein atoms being restrained each round (Supplementary Methods 1.1).

### Pose Generation with CHARMM Fast Fourier Transform Dock and Refinement

Docking grids representing protein and FAD atoms were generated in pyCHARMM^60^ with FFTG^19^, the CHARMM module for FFTDock, with a grid center at the average coordinates of 3MO in QM/MM refined TropB^27^ (Supplementary Methods 1.2). The top 500 poses were used as starting poses for grid-based minimization, explicit protein atom minimization, and simulated annealing. Grids using the same parameters as previously used for FFTDock^19^ were generated with varying epsilon to minimize the FFTDock poses (Supplementary Methods 1.3). For explicit protein minimization, AF2 structures with FAD cofactor were used to minimize the 500 FFTDock poses in vacuum with varying epsilon (Supplementary Methods 1.4). The top 10 poses from FFTDock were used as starting conformers to generate 500 total rotamers via random rotation and random translation for simulated annealing (Supplementary Methods 1.5). Simulated annealing was based on the CHARMM simulated annealing protocol using grids with varying softcore parameters and utilized CHARMM OpenMM_dock^19^ (OMMD) to carry out parallel simulated annealing of 500 rotamers.

### Stereochemistry Prediction

Each docking approach generated 500 final poses that were clustered using cluster.pl from the MMTSB toolset^61^, with parameters kclust, nolsqfit, 1 Å radius, and heavy atoms. For each cluster, the lowest energy pose was chosen as the cluster representative and the energy of this representative was used to rank the clusters. The stereochemistry of a cluster was calculated using the representative pose. From this pose, 3 atoms on the ligand resorcinol ring were selected to calculate a normal vector of the plane describing the ring. The normal vector was used to classify the representative pose as R or S based on its orientation relative to the vector from the ligand average coordinates to the FAD ring. The angle of a pose was constructed as the angle between the plane normal vector and the vector to the FAD ring. An angle between 0° and 90° indicated an S pose and an angle between 90° and 180° indicated an R pose. Each cluster was assigned a size based on the number of members of that cluster, a cluster energy determined by the representative pose, and a predicted stereochemistry and angle based on the representative pose (Figure S6).

The protein-ligand complex was assigned an overall predicted R fraction utilizing the average of the Boltzmann distribution^62^ of each cluster:

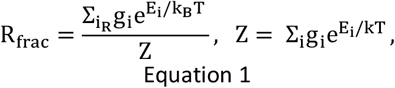

where i represents the cluster index, E_i_ is the energy of the representative cluster pose, k_B_ is the Boltzmann constant, g_i_ is the size of the cluster and the summation in the numerator is restricted to those clusters for which the geometry of the representative pose is identified to produce R stereochemistry and the summation in the denominator (Z) is over all clusters. Stereochemistry labels were assigned using an R_frac_ greater than 0.8 as R stereochemistry, an R_frac_ less that 0.2 as S stereochemistry, and racemic (R/S) for 0.2 < R_frac_ < 0.8. Accuracy of a docking approach was measured by comparing predicted stereochemistry labels against the ancestral FDMO library observations for the subset of reactive ligands. The average stereochemistry of a protein was the mode of the stereochemistry across all ligands, with R/S in case of multiple modes.

### Reactivity Prediction

A logistic regression model using statsmodels^63^ was used to fit the reactivity of an enzyme in the ancestral FDMO library as a binary classification problem. For training, an enzyme was labeled reactive if it showed any non-zero conversion with any ligand in the library. The logistic regression model considered five computed descriptors to predict reactivity: FAD distance, FAD angle, anion distance, docking energy efficiency, and Pafnucy^52^ predicted pK_a_ efficiency. Docking energy efficiency was found by taking the energy of the top ranked cluster pose and dividing it by the number of ligand heavy atoms. The Pafnucy predicted pK_d_ efficiency was found by dividing the predicted pK_d_ from Pafnucy for the complex with the protein, FAD, and the top ranked cluster representative pose by the number of ligand heavy atoms. FAD distance was defined as the distance from the ligand average coordinates of the top pose to the atom on the FAD ring that bonds to the hydroperoxyl group in the activated FAD. The FAD angle is the angle from the top cluster derived from stereochemistry prediction. The anion distance is the distance from the top ranked ligand pose (the ligands are all anions) and the CZ atom of R206 (TropB numbering). The logistic regression model was trained on the ancestral FDMO library (67 enzymes previously assayed with **2-5**) and used to predict the conversion for all 830 enzymes.

### Functional Screen

The finalized functional screen involved predicting the structure of an enzyme with AF2 and the consensus sequence hits, incorporation of FAD, pose generation with FFTDock, minimization of FFTDock poses with minimization in vacuum in the environment of a fixed explicit all atom protein and epsilon of 0.75, prediction of stereochemistry from pose geometry, and prediction of reactivity with the trained logistic regression model. This protocol was used to predict stereochemistry and reactivity for the entire sequence library.

### Sequence Alignment and Preprocessing

All extant and ancestral sequences were aligned using Clustal Omega^64^ with default parameters to construct an MSA as input to the sequence-function models. The MSA was trimmed to columns of interest defined by a set of binding site residues and second shell residues. For each of the 67 assayed enzymes in the ancestral FDMO library, a list of binding site residues was defined as the union of any residue with a heavy atom within 4.5 Å of any ligand (**2**-**5**) heavy atom in the top pose across the studied ligands. A list of second shell residues was defined as any residue with a heavy atom within 4.5 Å of any binding site residue. The union of all binding site and second shell residues across the 67 enzymes was taken to define residues of interest. Feature preprocessing for modeling was done by dropping MSA columns that were not in the residues of interest or contained more than 10% gaps.

### Sequence-Function Model and SHAP Analysis

The mljar automated machine learning (AutoML) framework^54^ was used to train multiple classification models to predict average stereochemistry or consensus reactivity from the functional screen using the processed MSA. The input was the MSA with amino acid labels and transformation to numerical features was performed by mljar. Stereochemistry labels were defined as -1 (S), 0 (R/S), and 1 (R) from averaging the cutoffs applied to predicted R_frac_ for ligands **2-5**. Reactivity labels were defined as 0 (unreactive) or 1 (reactive) from the consensus logistic regression model using averaged reactivity descriptors for ligands **2-5**. For a comparison of all default models, we fitted an mljar AutoML using Explain mode, explain level 2, and algorithms: Baseline, Decision Tree^57,65^, Linear^57^, XGBoost^56^, Random Forest^57,66^, LightGBM^55,67^, CatBoost^55^, Neural Network^68^, and Nearest Neighbors^57^. Hyperparameter tuning was done using mljar AutoML in perform mode, explain level 2, no golden features, and CatBoost, XGBoost, and Random Forest algorithms. To rank the key residues, the mean absolute SHAP^42^ importance was normalized between 0 and 1 for each fold of each trained model. Then the normalized mean absolute SHAP importances were averaged across all generated models and their respective folds, to get a mean absolute SHAP importance of every residue across all trained models. To infer the effect of a single residue on stereochemistry or reactivity, a SHAP explainer^42,69^ was fitted to each fold of each model. The fold level SHAP values from the SHAP explainer were normalized between -1 and 1. For each residue, the normalized fold SHAP values were separated by amino acid type and plotted across all folds of all models to create a consensus dependence plot.

## Data Availability

All protein sequences, predicted structures, docked ligand poses, and predictions required to reproduce this study are available in the supplementary data (si_data) directory at https://github.com/BrooksResearchGroup-UM/seq_struct_func.

## Code Availability

Scripts to perform sequence-structure-function pipeline calculations and reproduce results using supplementary data are available at https://github.com/BrooksResearchGroup-UM/seq_struct_func.

## References

1. Brannigan, J. A. & Wilkinson, A. J. Protein engineering 20 years on. Nature Reviews Molecular Cell Biology 2002 3:12 3, 964–970 (2002).

2. Tsuboyama, K. et al.. Mega-scale experimental analysis of protein folding stability in biology and protein design. bioRxiv 2022.12.06.519132 (2022) doi:10.1101/2022.12.06.519132.

3. Packer, M. S. & Liu, D. R. Methods for the directed evolution of proteins. Nat Rev Genet. 16, 379–394 (2015).

4. Lutz, S. Beyond directed evolution-semi-rational protein engineering and design. Curr Opin Biotechnol. 21, 734–743 (2010).

5. Song, Z., Zhang, Q., Wu, W., Pu, Z. & Yu, H. Rational design of enzyme activity and enantioselectivity. Front Bioeng Biotechnol 11, 91 (2023).

6. Jumper, J. et al.. Highly accurate protein structure prediction with AlphaFold. Nature 2021 596:7873 596, 583–589 (2021).

7. Jumper, J. et al.. Applying and improving AlphaFold at CASP14. Proteins: Structure, Function, and Bioinformatics 89, 1711–1721 (2021).

8. Varadi, M. et al.. AlphaFold Protein Structure Database: massively expanding the structural coverage of protein-sequence space with high-accuracy models. Nucleic Acids Res 50, D439–D444 (2022).

9. Lin, Z. et al.. Evolutionary-scale prediction of atomic-level protein structure with a language model. Science (1979) 379, 1123–1130 (2023).

10. Bouatta, N. & AlQuraishi, M. Structural biology at the scale of proteomes. Nat Struct Mol Biol 30, 129–130 (2023).

11. AlQuraishi, M. End-to-End Differentiable Learning of Protein Structure. Cell Syst 8, 292-301.e3 (2019).

12. Mirdita, M. et al.. ColabFold: making protein folding accessible to all. Nature Methods 2022 19:6 19, 679–682 (2022).

13. Kandathil, S. M., Greener, J. G., Lau, A. M. & Jones, D. T. Ultrafast end-to-end protein structure prediction enables high-throughput exploration of uncharacterized proteins. Proc Natl Acad Sci U S A 119, e2113348119 (2022).

14. Baek, M. et al.. Accurate prediction of protein structures and interactions using a three-track neural network. Science 373, 871–876 (2021).

15. Mirdita, M., Steinegger, M. & Söding, J. MMseqs2 desktop and local web server app for fast, interactive sequence searches. Bioinformatics 35, 2856–2858 (2019).

16. Lee, D., Redfern, O. & Orengo, C. Predicting protein function from sequence and structure. Nat Rev Mol Cell Biol (2007) doi:10.1038/nrm2281.

17. Perkins, R., Fang, H., Tong, W. & Welsh, W. J. Quantitative structure-activity relationship methods: perspectives on drug discovery and toxicology. Environ Toxicol Chem 22, 1666–1679 (2003).

18. Meng, X.-Y., Zhang, H.-X., Mezei, M. & Cui, M. Molecular Docking: A powerful approach for structure-based drug discovery. Curr Comput Aided Drug Des 7, 146 (2011).

19. Ding, X., Wu, Y., Wang, Y., Vilseck, J. Z. & Brooks C. L. III. Accelerated CDOCKER with GPUs, Parallel Simulated Annealing, and Fast Fourier Transforms. J Chem Theory Comput 16, 3910–3919 (2020).

20. Brooks, B. R. et al.. CHARMM: The Biomolecular Simulation Program. J Comput Chem 30, 1545 (2009).

21. Huang, J. et al.. CHARMM36m: an improved force field for folded and intrinsically disordered proteins. Nat Methods 14, 71–73 (2017).

22. Vanommeslaeghe, K. et al.. CHARMM General Force Field (CGenFF): A force field for drug-like molecules compatible with the CHARMM all-atom additive biological force fields. J Comput Chem 31, 671 (2010).

23. Baker Dockrey, S. A., Lukowski, A. L., Becker, M. R. & Narayan, A. R. H. Biocatalytic site- and enantioselective oxidative dearomatization of phenols. Nature Chemistry 2017 10:2 10, 119–125 (2017).

24. Pyser, J. B. et al.. Stereodivergent, chemoenzymatic synthesis of azaphilone natural products. J Am Chem Soc 141, 18551 (2019).

25. Tweedy, S. E. et al.. Hydroxyl Radical-Coupled Electron-Transfer Mechanism of Flavin-Dependent Hydroxylases. Journal of Physical Chemistry B 123, 8065–8073 (2019).

26. Baker Dockrey, S. A. et al. Positioning-Group-Enabled Biocatalytic Oxidative Dearomatization. ACS Cent. Sci 5, 1010–1016 (2019).

27. Rodríguez Benítez, A. et al.. Structural Basis for Selectivity in Flavin-Dependent Monooxygenase-Catalyzed Oxidative Dearomatization. ACS Catal 9, 3633–3640 (2019).

28. Nicoll, C. R. et al.. Ancestral-sequence reconstruction unveils the structural basis of function in mammalian FMOs. Nature Structural & Molecular Biology 2019 27:1 27, 14–24 (2019).

29. Chiang, C.-H. et al.. Deciphering the evolution of flavin-dependent monooxygenase stereoselectivity using ancestral sequence reconstruction. Proceedings of the National Academy of Sciences 120, e2218248120 (2023).

30. Aadland, K., Pugh, C. & Kolaczkowski, B. High-Throughput Reconstruction of Ancestral Protein Sequence, Structure, and Molecular Function. Methods in Molecular Biology 1851, 135–170 (2019).

31. Eswar, N. et al.. Comparative protein structure modeling using Modeller. Curr Protoc Bioinformatics Chapter 5, (2006).

32. Wong, F. et al.. Benchmarking AlphaFold-enabled molecular docking predictions for antibiotic discovery. Mol Syst Biol 18, e11081 (2022).

33. Eberhardt, J., Santos-Martins, D., Tillack, A. F. & Forli, S. AutoDock Vina 1.2.0: New Docking Methods, Expanded Force Field, and Python Bindings. J. Chem. Inf. Model 61, 3891–3898 (2021).

34. Wijma, H. J. et al.. Enantioselective enzymes by computational design and in silico screening. Angewandte Chemie - International Edition 54, 3726–3730 (2015).

35. Wijma, H. J., Marrink, S. J. & Janssen, D. B. Computationally Efficient and Accurate Enantioselectivity Modeling by Clusters of Molecular Dynamics Simulations. J. Chem. Inf. Model. 54, 2079–2092 (2014).

36. Arabnejad, H. et al.. Computational Design of Enantiocomplementary Epoxide Hydrolases for Asymmetric Synthesis of Aliphatic and Aromatic Diols. ChemBioChem 21, 1893–1904 (2020).

37. Hekkelman, M. L., de Vries, I., Joosten, R. P. & Perrakis, A. AlphaFill: enriching AlphaFold models with ligands and cofactors. Nat Methods (2022) doi:10.1038/s41592-022-01685-y.

38. Krieger, E., Koraimann, G. & Vriend, G. Increasing the precision of comparative models with YASARA NOVA—a self-parameterizing force field. Proteins: Structure, Function, and Bioinformatics 47, 393–402 (2002).

39. Yang, K. K., Wu, Z. & Arnold, F. H. Machine-learning-guided directed evolution for protein engineering. Nat Methods 16, 687–694 (2019).

40. Wu, Z., Jennifer Kan, S. B., Lewis, R. D., Wittmann, B. J. & Arnold, F. H. Machine learning-assisted directed protein evolution with combinatorial libraries. Proc Natl Acad Sci U S A 116, 8852–8858 (2019).

41. Cadet, F. et al.. A machine learning approach for reliable prediction of amino acid interactions and its application in the directed evolution of enantioselective enzymes. Scientific Reports 2018 8:1 8, 1–15 (2018).

42. Lundberg, S. M., Allen, P. G. & Lee, S.-I. A Unified Approach to Interpreting Model Predictions. in Advances in Neural Information Processing Systems 30 4765–4774 (2017).

43. Linardatos, P., Papastefanopoulos, V. & Kotsiantis, S. Explainable AI: A Review of Machine Learning Interpretability Methods. Entropy 2021, Vol. 23, Page 18 23, 18 (2020).

44. Bi, Y. et al.. An Interpretable Prediction Model for Identifying N7-Methylguanosine Sites Based on XGBoost and SHAP. Mol Ther Nucleic Acids 22, 362–372 (2020).

45. Rodríguez-Pérez, R. & Bajorath, J. Interpretation of machine learning models using shapley values: application to compound potency and multi-target activity predictions. J Comput Aided Mol Des 34, 1013–1026 (2020).

46. Zhang, Y. & Skolnick, J. TM-align: a protein structure alignment algorithm based on the TM-score. Nucleic Acids Res 33, 2302–2309 (2005).

47. Case, D. A. et al.. The Amber Biomolecular Simulation Programs. J Comput Chem 26, 1668–1688 (2005).

48. Mariani, V., Biasini, M., Barbato, A. & Schwede, T. lDDT: a local superposition-free score for comparing protein structures and models using distance difference tests. Bioinformatics 29, 2722–2728 (2013).

49. Wang, Z. et al.. Comprehensive evaluation of ten docking programs on a diverse set of protein–ligand complexes: the prediction accuracy of sampling power and scoring power. Physical Chemistry Chemical Physics 18, 12964–12975 (2016).

50. Baldi, P., Brunak, S., Chauvin, Y., Andersen, C. A. F. & Nielsen, H. Assessing the accuracy of prediction algorithms for classification: an overview. Bioinformatics 16, 412–424 (2000).

51. Gorodkin, J. Comparing two K-category assignments by a K-category correlation coefficient. Comput Biol Chem 28, 367–374 (2004).

52. Stepniewska-Dziubinska, M. M., Zielenkiewicz, P. & Siedlecki, P. Development and evaluation of a deep learning model for protein-ligand binding affinity prediction. Bioinformatics 34, 3666–3674 (2018).

53. Wang, R., Fang, X., Lu, Y., Yang, C.-Y. & Wang, S. The PDBbind Database: Methodologies and Updates. (2005) doi:10.1021/jm048957q.

54. Plonska, A. & Plonski, P. MLJAR: State-of-the-art Automated Machine Learning Framework for Tabular Data.. Preprint at https://github.com/mljar/mljar-supervised (2021).

55. Dorogush, A. V., Ershov, V., Gulin, A. & Yandex. CatBoost: gradient boosting with categorical features support. 1810.11363 [cs.LG] (2018) doi:10.48550/arXiv.1810.11363.

56. Chen, T. & Guestrin, C. XGBoost: A Scalable Tree Boosting System. 1603.02754 [cs.LG] (2016) doi:10.48550/arXiv.1603.02754.

57. Pedregosa, F. et al.. Scikit-learn: Machine Learning in Python. Journal of Machine Learning Research 12, 2825–2830 (2011).

58. Steinegger, M. et al.. HH-suite3 for fast remote homology detection and deep protein annotation. BMC Bioinformatics 20, 1–15 (2019).

59. Zhang, Y. & Skolnick, J. TM-align: a protein structure alignment algorithm based on the TM-score. doi:10.1093/nar/gki524.

60. Buckner, J. et al.. pyCHARMM: Embedding CHARMM Functionality in a Python Framework. J Chem Theory Comput 19, 3752–3762 (2023).

61. Feig, M., Karanicolas, J. & Brooks C. L. III. MMTSB Tool Set: enhanced sampling and multiscale modeling methods for applications in structural biology. J Mol Graph Model 22, 377–395 (2004).

62. Gibbs, J. W. Elementary Principles in Statistical Mechanics: Developed with Especial Reference to the Rational Foundation of Thermodynamics. Cambridge Library Collection - Mathematics (Cambridge University Press, 2010). doi:10.1017/CBO9780511686948.

63. Seabold, S. & Perktold, J. Statsmodels: Econometric and statistical modeling with Python. in 9th Python in Science Conference (2010).

64. Sievers, F. et al.. Fast, scalable generation of high-quality protein multiple sequence alignments using Clustal Omega. Mol Syst Biol 7, 539 (2011).

65. Breiman, L. Classification and regression trees. (Routledge, 2017).

66. Breiman, L. Random Forests. Mach Learn 45, 5–32 (2001).

67. Ke, G. et al.. LightGBM: A Highly Efficient Gradient Boosting Decision Tree. in Advances in Neural Information Processing Systems 30 3149–3157 (2017).

68. Abadi, M. et al.. Tensorflow: a system for large-scale machine learning. in Osdi vol. 16 265–283 (Savannah, GA, USA, 2016).

69. Lundberg, S. M. et al.. From local explanations to global understanding with explainable AI for trees. Nature Machine Intelligence 2020 2:1 2, 56–67 (2020).

