## Supplementary Materials for "Guiding Discovery of Protein Sequence-Structure-Function Modeling"

Azam Hussain[a] and Charles L. Brooks III[a,b,c]

[a]Macromolecular Science and Engineering Program, University of Michigan, Ann Arbor, MI 48109, United States
[b]Department of Chemistry, University of Michigan, Ann Arbor, MI 48109, United States
[c]Biophysics Program, University of Michigan, Ann Arbor, MI 48109, United States

### Supplementary Methods 1

#### **1.1** Minimization and Addition of FAD Cofactor

The AF2 models were superposed with TM-align[1] with to a previously generated QM/MM refined chain A of RCSB PDB 6NES docked with 3-methyl-orcinaldeyde (3MO)[2]. The MMTSB tool convpdb.pl[3] was used to reformat the models into the CHARMM[4] PDB-consistent format. The superposed structures were represented using in CHARMM in vacuum using the CHARMM36 forcefield[5] and a constant dielectric (CDIE) with epsilon 1 using the python package pyCHARMM[6]. The non-bonded switching and interaction cutoffs for van der Waals and electrostatic interactions were FSWITCH, VFSWITCH, CUTNB of 10 Å, CTOFNB of 8 Å, and CTONNB of 7 Å. The superposed structures were minimized with harmonic restraints using a force constant of 50 kcal/mol/Å$^2$ on all heavy atoms, using 1000 steps of steepest descents (SD) (TOLENR of .001 and TOLGRD of .0001). Next, the FAD cofactor present in the QM/MM structure was added to the models. The model structure with FAD had harmonic restraints placed on all protein backbone heavy atoms with a force constant of 50 kcal/mol/Å$^2$ and was minimized with 200 steps of SD (TOLENR of .001 and TOLGRD of .0001). Following, further minimization (200 steps SD, TOLENR of .001, TOLGRD of .0001) with harmonic restraints using a force constant of 50 kcal/mol/Å$^2$ on all side chain heavy atoms further than 5 Å from FAD was performed. Finally, all heavy atoms except FAD were restrained using harmonic restraints and minimized (force constant 50 kcal/mol/Å$^2$, 200 steps of SD followed by 1000 steps of Adopted Basis Newton Raphson (ABNR) minimization with TOLENR of .001, TOLGRD of .0001).

#### **1.2** Pose Generation with CHARMM Fast Fourier Transform Dock

Ligand protomers were generated with MOE v.2019.0102 (Chemical Computing Group, Montreal, Canada) and top protomers consistent with the TropB native substrate previously used[2] were selected. Docking grids representing protein and FAD atoms were generated in pyCHARMM[6] with CHARMM FFTG[7], using a grid spacing of 0.5 Å, grid center at the average coordinates of 3MO in QM/MM refined TropB[2], grid dimensions of 12Å × 12Å ×12Å, softcore potentials using EMAX of 2 kcal/mol, MINE of -20 kcal/mol and MAXE of 40 kcal/mol, distance-dependent dielectric constant (RDIE) with an epsilon of 3, CUTNB of 999, CTOFNB of 999, and CTONNB of 999. Ligands were parameterized using CGenFF[8]. Ligands were docked to the grids with CHARMM FFTG[7] using 36000 saved poses, a set of 36,863 quaternions, and SIZB of 100. The top 500 poses were selected with the difference between the ligand pose grid energy and initial ligand energy in vacuum reported. The poses were used as starting poses for grid-based minimization, explicit protein atom minimization, and simulated annealing.

**1.3** Grid-based Minimization

Grids using the same parameters as previously used for FFTDock[7] were generated with varying epsilon. The top 500 poses from FFTDock were minimized using 50 steps of SD and 1000 steps of ABNR with a TOLENR of .1. The reported docking energy was the final minimized grid energy subtracted by the initial energy of the ligand in vacuum.

**1.4** Explicit Atom Minimization

Protein structures with FAD cofactor were used to minimize the 500 FFTDock[7] poses in vacuum, with CUTNB of 12 Å, CTOFNB of 10 Å, CTONNB of 10 Å, RDIE, varying epsilon, SWITCH, and VSWITCH. Protein and FAD atoms were fixed with CONS FIX. Minimization was performed with 50 steps of SD followed by 1000 steps of ABNR with TOLENR of .001. The reported docking energy was the final minimized ligand energy subtracted by the initial ligand energy.

**1.5** Simulated Annealing

Docking based on the CHARMM simulated annealing protocol[7] was performed in vacuum with SWITCH, VSWITCH, CUTNB of 12 Å, CTOFNB of 10 Å, and CTONNB of 8 Å, and RDIE with varying epsilon. Grids were generated with the same protocol for FFTDock but varying softcore parameters. The soft grid used an EMAX of 3 kcal/mol, MINE of -20 kcal/mol, and MAXE of 40 kcal/mol. The mine grid used an EMAX of 15 kcal/mol, MINE of -120 kcal/mol, and MAXE of -2 kcal/mol. The hard grid used an EMAX of 100 kcal/mol, MINE of -100 kcal/mol, and MAXE of 100 kcal/mol. CHARMM OpenMM_dock[7] (OMMD) was used to carry out parallel simulated annealing on the 500 rotamers. OMMD grid was used to read in the mine grid and soft grid. The top 10 poses from FFTDock were used as starting conformers to generate 500 total rotamers for simulated annealing. The ligand selection and NUMCOPY of 500 were passed to OMMD build. Starting conformers were briefly minimized in vacuum with 50 steps of SD with TOLENR of 0.01 and 50 steps of ABNR with TOLENR of 0.005. 50 rotamers were generated for each conformer pose for a total of 500 poses. Each rotamer was generated by random rotation of the starting conformer between 0 and 180 degrees, random translation between 0 and 2 Å, minimization with 100 steps of SD and 500 steps of ABNR with TOLENR of 0.01 in the soft grid, and lastly minimization in the hard grid using 50 steps of SD and 100 steps of conjugate gradient descent (CONJ) with TOLENR of 0.01. OMMD simulated annealing (SIAN) was carried out on the 500 rotamers with soft potentials, 3000 steps, starting temperature of 300 K, final temperature of 700 K, NHRQ 50, and INCT 1. Next, SIAN was carried out with soft potential, 14000 steps, starting temperature of 700K and final temperature of 300 K. SIAN was then carried out with hard potential, 7000 steps, starting temperature of 500 K and ending temperature of 300 K. Lastly, SIAN was performed with hard potential, 300 steps, starting temperature of 400K, final temperature of 50K. Final poses were minimized with 50 steps of SD and 1000 steps of ABNR and TOLENR of 0.001.

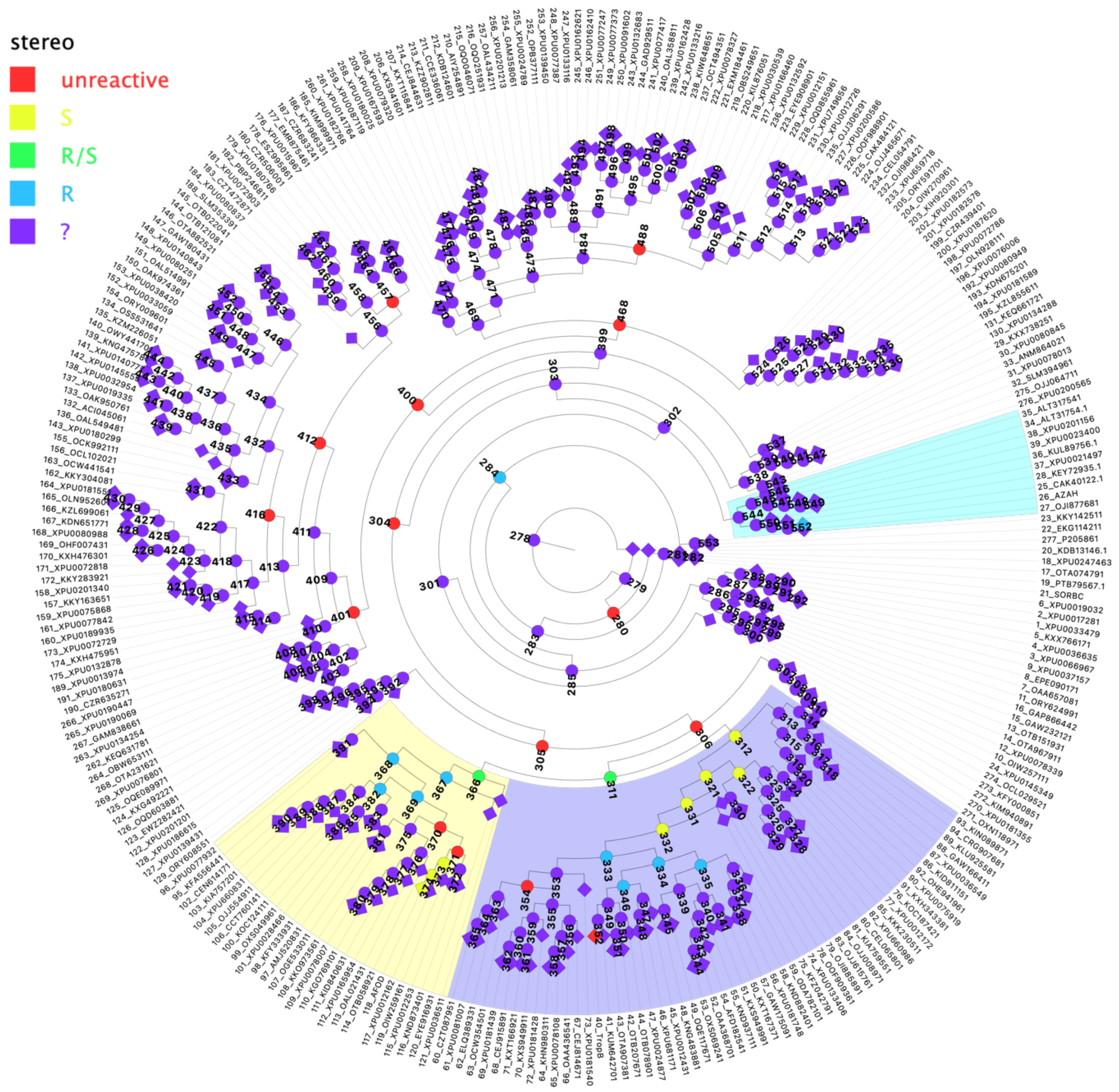

Figure S1: Phylogenetic tree displaying scope of ancestral FDMO library[9]. Ancestors are represented as spheres extant sequences are represented as diamonds. Based on the preliminary screen of 67 members from this tree, unreactive sequences are shown as red, yellow denotes S stereochemistry, green racemic, and blue denotes R stereochemistry. Purple represents sequences for which the functional properties are unknown because they have not been expressed or tested. Known protein clades are indicated by the shaded background with blue corresponding to the TropB clade, yellow the AfoD clade, and cyan is the AzaH clade.

Table S1: Comparison of different protein structure prediction methods for TropB. AlphaFold2_cons indicates using MSA generation preconditioned on consensus sequence hits with AlphaFold2 pipeline. AlphaFold2_dummy indicates using dummy templates instead of PDB template hits.

| **Model** | **RMSD** | **TM-score** | **GDT-TS** | **GDT-TS** |
|---|---|---|---|---|
| **Alphafold2**[10] | 0.808 | 0.9574 | 0.9541 | 0.9541 |
| **Alphafold2_cons** | 0.91 | 0.9573 | 0.953 | 0.953 |
| **Alphafold2_dummy** | 1.561 | 0.9339 | 0.8507 | 0.8507 |
| **Alphafold2_cons_dummy** | 1.528 | 0.9401 | 0.8708 | 0.8708 |
| **I-TASSER**[11] | 0.194 | 0.9636 | 0.9642 | 0.9642 |
| **RaptorX**[12] | 6.717 | 0.8036 | 0.6102 | 0.6102 |
| **RosettaCM**[13] | 0.763 | 0.957 | 0.953 | 0.953 |
| **RoseTTAfold**[14] | 2.096 | 0.9109 | 0.7886 | 0.7886 |
| **SwissModel**[15] | 0.065 | 0.9862 | 0.9863 | 0.9863 |

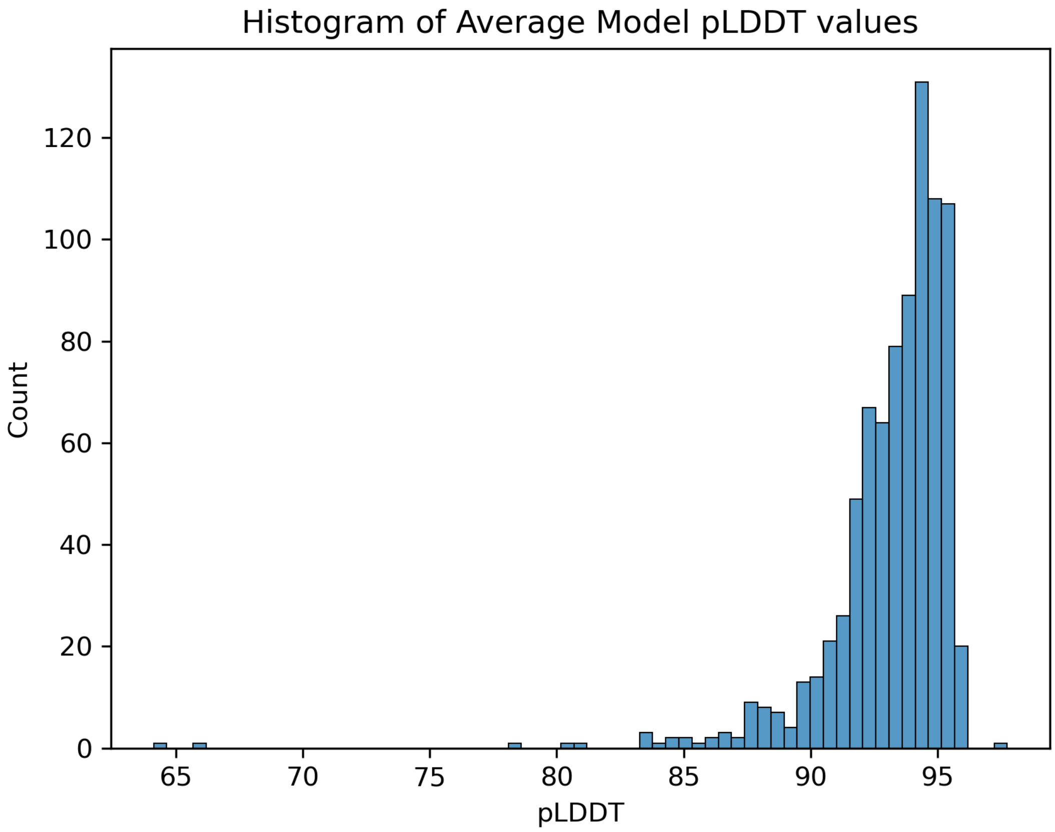

Figure S2: Histogram of average pLDDT[16] values across models. pLDDT values were averaged across all residues.

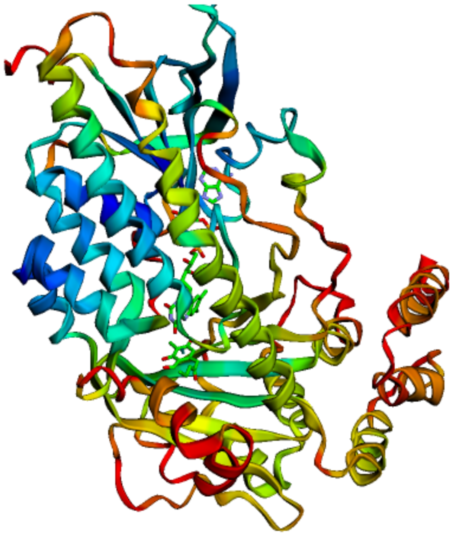

Figure S3: Anc 278 colored by predicted Local Distance Test (pLDDT[16]) from AlphaFold2[10]. Red indicates low scoring regions and blue indicates high scoring regions. The FAD cofactor and docked ligand represent the binding site which posess more conserved residues.

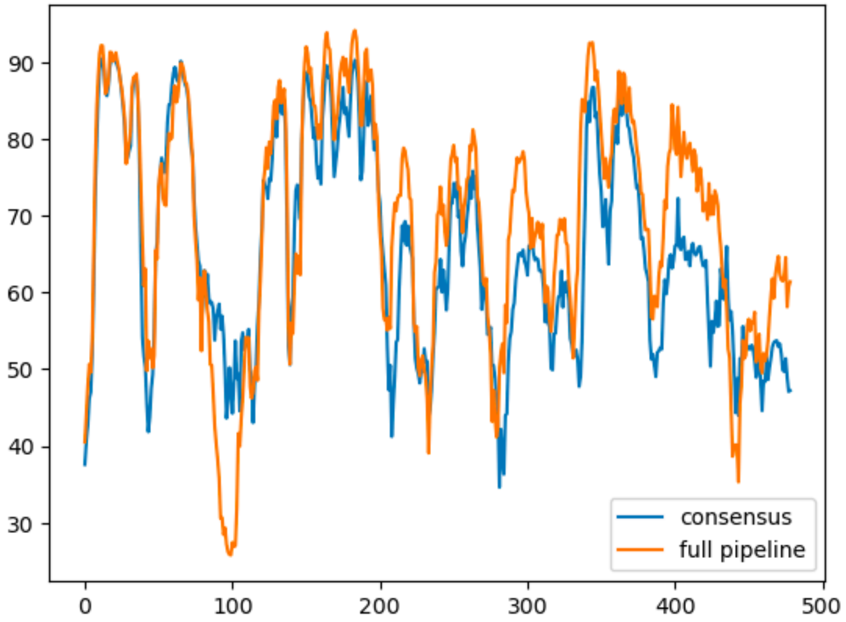

Figure S4: Anc 278 pLDDT[16] per residue scores for AlphaFold2[10] pipeline conditioned on consensus MSA versus full AlphaFold pipeline. Both pipelines used monomer model 1 and default parameters. The mean pLDDT score for the consensus model was 65.7 and the mean pLDDT score for the full pipeline was 69.3. The $C\alpha$ RMSD using TM-align[1] was 2.88 Å between the consensus pipeline and full pipeline Anc 278 structures.

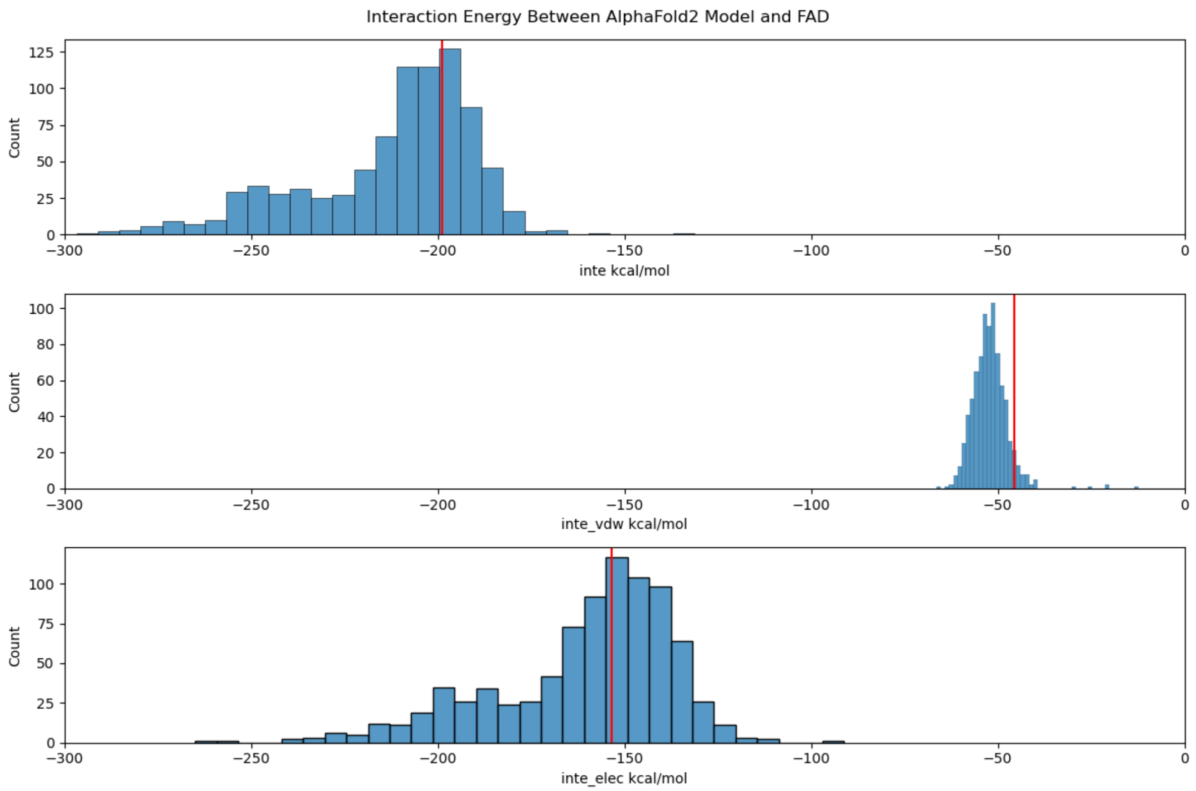

Figure S5: Interaction energy between AlphaFold2 structures and FAD cofactor. Red line indicates energy of TropB. Top panel is total interaction energy in kcal/mol, middle panel depicts van der Waals hydrophobic energy, and the bottom panel electorstatic interaction energy.

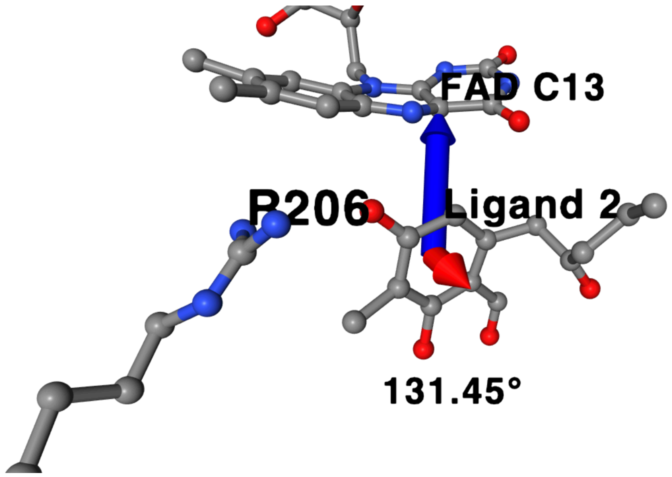

Figure S6: Prediction of stereochemistry from docked structures. The red vector indicates the normal vector the plane of the ligand resorcinol ring defined by three atoms. The blue vector indicates the vector from the ligand resorcinol ring and anion average coordinates to the FAD C13 atom. The angle between the blue and red vectors (131.45°) is used to classify the stereochemistry of a pose as R or S, with an angle greater than 90° as R, otherwise as S.

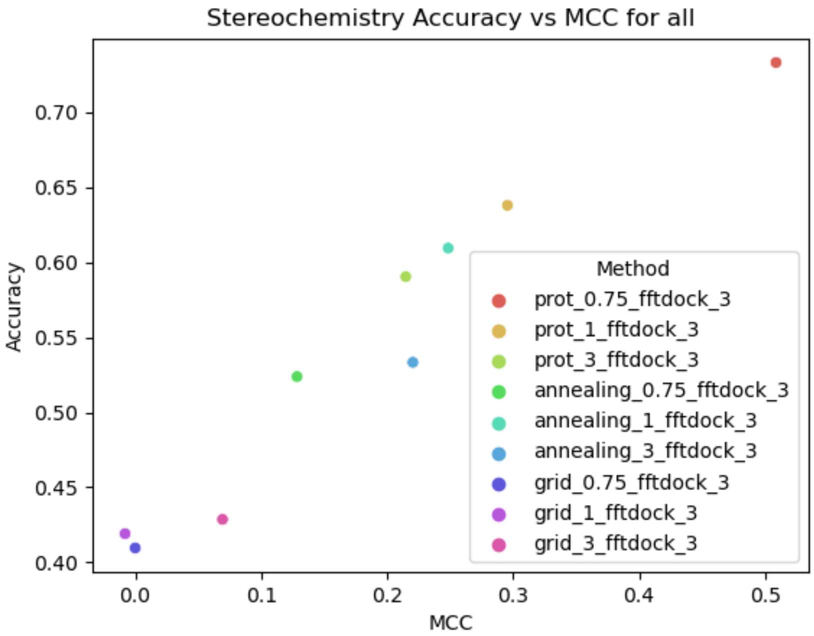

Figure S7: Docking methods MCC vs Accuracy for all protein ligand stereochemistry pairs. "Prot" is minimization of FFTDock[7] poses in explicit protein. "Annealing" is use of FFTDock poses as starting poses in simulated annealing. "Grid" is minimization of FFTDock poses further in soft grid potentials. The rescoring approaches are named as follows: (method)_(method epsilon)_fftdock_(fftdock epsilon). All methods used FFTDock poses generated at epsilon of 3.

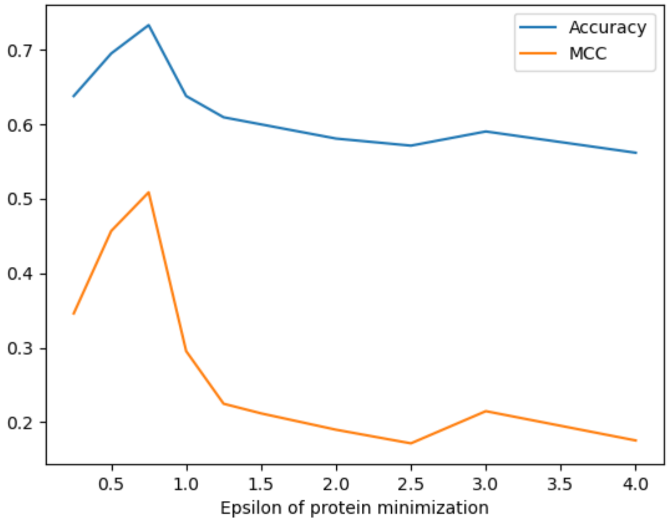

Figure S8: Accuracy and MCC for stereochemistry prediction against varying epsilon for protein minimization. Accuracy and MCC were calculated using all protein ligand pairs with experimental stereochemistry data. Protein minimization was performed by minimizing FFTDock[7] poses in explicit protein, with FFTDock poses generated at epsilon of 3.

Table S2: Stereochemistry classification scores for various docking approaches (-1.0: S, 1.0: R)

| Method | Ligand | MCC | Accuracy | -1.0 precision | -1.0 recall | -1.0 support | 1.0 precision | 1.0 recall | 1.0 support |
|---|---|---|---|---|---|---|---|---|---|
| prot_0.75 | 18-2 | 0.73 | 0.85 | 1.00 | 0.67 | 15.00 | 0.79 | 1.00 | 19.00 |
| | 17-2 | 0.35 | 0.61 | 0.46 | 0.86 | 7.00 | 0.89 | 0.50 | 16.00 |
| | ome | 0.68 | 0.81 | 0.73 | 1.00 | 8.00 | 1.00 | 0.69 | 13.00 |
| | ketone | 0.43 | 0.63 | 0.57 | 1.00 | 12.00 | 1.00 | 0.33 | 15.00 |
| | all | 0.51 | 0.73 | 0.65 | 0.86 | 42.00 | 0.87 | 0.65 | 63.00 |
| prot_1 | 18-2 | 0.52 | 0.74 | 1.00 | 0.40 | 15.00 | 0.68 | 1.00 | 19.00 |
| | 17-2 | 0.19 | 0.52 | 0.42 | 0.71 | 7.00 | 0.78 | 0.44 | 16.00 |
| | ome | 0.36 | 0.67 | 0.55 | 0.75 | 8.00 | 0.80 | 0.62 | 13.00 |
| | ketone | 0.31 | 0.59 | 0.55 | 0.92 | 12.00 | 0.83 | 0.33 | 15.00 |
| | all | 0.30 | 0.64 | 0.57 | 0.67 | 42.00 | 0.74 | 0.62 | 63.00 |
| prot_3 | 18-2 | 0.17 | 0.59 | 0.75 | 0.20 | 15.00 | 0.59 | 0.89 | 19.00 |
| | 17-2 | 0.44 | 0.74 | 0.56 | 0.71 | 7.00 | 0.86 | 0.75 | 16.00 |
| | ome | 0.05 | 0.52 | 0.38 | 0.38 | 8.00 | 0.67 | 0.62 | 13.00 |
| | ketone | 0.33 | 0.52 | 0.61 | 0.92 | 12.00 | 1.00 | 0.20 | 15.00 |
| | all | 0.21 | 0.59 | 0.56 | 0.52 | 42.00 | 0.69 | 0.63 | 63.00 |
| annealing_0.75 | 18-2 | 0.00 | 0.53 | 0.44 | 0.27 | 15.00 | 0.56 | 0.74 | 19.00 |
| | 17-2 | 0.23 | 0.52 | 0.38 | 0.86 | 7.00 | 0.86 | 0.38 | 16.00 |
| | ome | 0.34 | 0.52 | 0.50 | 1.00 | 8.00 | 1.00 | 0.23 | 13.00 |
| | ketone | 0.20 | 0.52 | 0.56 | 0.83 | 12.00 | 0.67 | 0.27 | 15.00 |
| | all | 0.13 | 0.52 | 0.47 | 0.67 | 42.00 | 0.66 | 0.43 | 63.00 |
| annealing_1 | 18-2 | 0.10 | 0.56 | 0.50 | 0.27 | 15.00 | 0.60 | 0.79 | 19.00 |
| | 17-2 | 0.09 | 0.48 | 0.33 | 0.71 | 7.00 | 0.75 | 0.38 | 16.00 |
| | ome | 0.41 | 0.67 | 0.54 | 0.88 | 8.00 | 0.88 | 0.54 | 13.00 |
| | ketone | 0.58 | 0.74 | 0.67 | 1.00 | 12.00 | 1.00 | 0.53 | 15.00 |
| | all | 0.25 | 0.61 | 0.52 | 0.67 | 42.00 | 0.73 | 0.57 | 63.00 |
| annealing_3 | 18-2 | 0.30 | 0.62 | 0.60 | 0.40 | 15.00 | 0.75 | 0.79 | 19.00 |
| | 17-2 | 0.10 | 0.43 | 0.31 | 0.71 | 7.00 | 0.83 | 0.31 | 16.00 |
| | ome | 0.29 | 0.52 | 0.47 | 0.88 | 8.00 | 1.00 | 0.31 | 13.00 |
| | ketone | 0.29 | 0.52 | 0.56 | 0.75 | 12.00 | 1.00 | 0.33 | 15.00 |
| | all | 0.22 | 0.53 | 0.47 | 0.64 | 42.00 | 0.83 | 0.46 | 63.00 |
| grid_0.75 | 18-2 | -0.04 | 0.41 | 0.45 | 0.87 | 15.00 | 0.33 | 0.05 | 19.00 |
| | 17-2 | -0.20 | 0.26 | 0.24 | 0.57 | 7.00 | 0.50 | 0.13 | 16.00 |
| | ome | 0.20 | 0.52 | 0.50 | 0.75 | 8.00 | 0.71 | 0.38 | 13.00 |
| | ketone | -0.01 | 0.44 | 0.43 | 0.75 | 12.00 | 0.60 | 0.20 | 15.00 |
| | all | 0.00 | 0.41 | 0.41 | 0.76 | 42.00 | 0.58 | 0.17 | 63.00 |
| grid_1 | 18-2 | 0.04 | 0.47 | 0.46 | 0.80 | 15.00 | 0.57 | 0.21 | 19.00 |
| | 17-2 | -0.47 | 0.17 | 0.07 | 0.14 | 7.00 | 0.43 | 0.19 | 16.00 |
| | ome | 0.12 | 0.43 | 0.44 | 0.88 | 8.00 | 0.67 | 0.15 | 13.00 |
| | ketone | 0.21 | 0.56 | 0.50 | 0.67 | 12.00 | 0.78 | 0.47 | 15.00 |
| | all | -0.01 | 0.42 | 0.39 | 0.67 | 42.00 | 0.62 | 0.25 | 63.00 |
| grid_3 | 18-2 | -0.14 | 0.29 | 0.38 | 0.60 | 15.00 | 0.33 | 0.05 | 19.00 |
| | 17-2 | 0.10 | 0.61 | 0.00 | 0.00 | 7.00 | 0.74 | 0.88 | 16.00 |
| | ome | 0.19 | 0.43 | 0.42 | 1.00 | 8.00 | 1.00 | 0.08 | 13.00 |
| | ketone | 0.09 | 0.44 | 0.50 | 0.75 | 12.00 | 0.60 | 0.20 | 15.00 |
| | all | 0.07 | 0.43 | 0.43 | 0.62 | 42.00 | 0.68 | 0.30 | 63.00 |

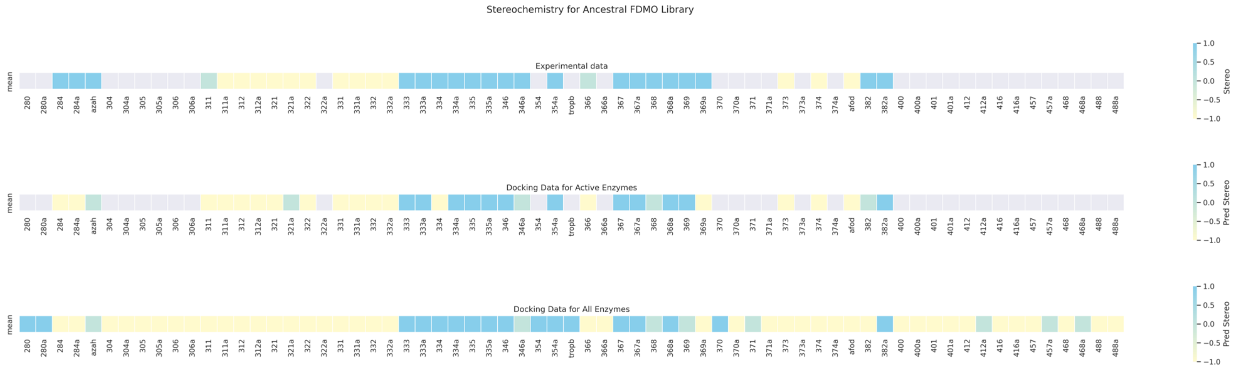

Figure S9: Mean stereochemistry results for ancestral FDMO library. Blue squares represent R stereochemistry, yellow squares represent S stereochemistry, and greens squares represent racemic stereochemistry. The top panel indicates mean stereochemistry for previous experimental results. The middle panel indicates mean stereochemistry for protocol prediction for active enzymes. The bottom panel indicates mean stereochemistry for protocol predictions for all enzymes in ancestral FDMO library.

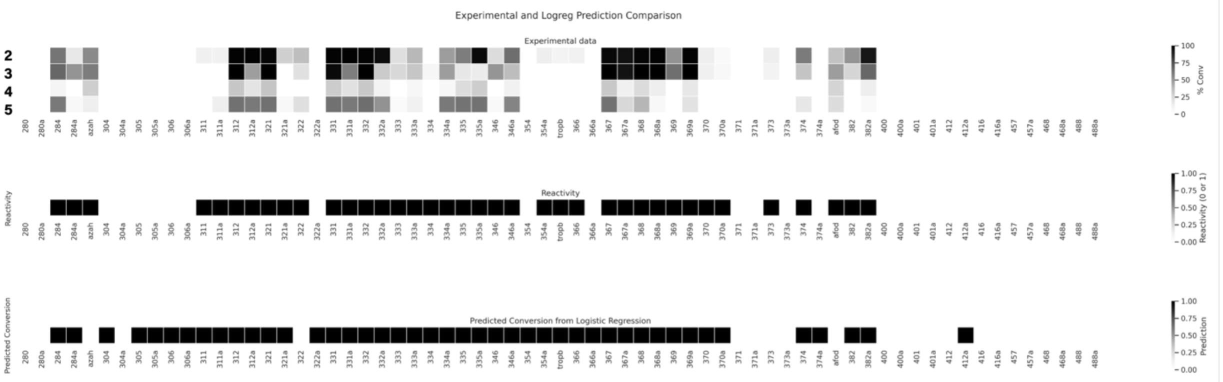

Figure S10: Predicted reactivity from logistic regression versus experimental conversion for the ancestral FDMO library. The top panel is the original conversion data for the experimental assay, with black indicating conversion near 100, and white representing 0 conversion. The middle panel is the consensus reactivity label assigned by any nonzero conversion with any ligand. The bottom panel is the predictions from consensus logistic regression model.

Table S3: Logistic regression model parameters. Average Pafnucy[17] p$K_d$ efficiency is an order of magnitude smaller than average docking energy efficiency, therefore coefficients between the two approaches are comparable.

| | | | | | | |
|---|---|---|---|---|---|---|
| **Dep. Variable:** | Conversion | **No. Observations:** | 67 | | | |
| **Model:** | Logit | **Df Residuals:** | 63 | | | |
| **Method:** | MLE | **Df Model:** | 3 | | | |
| **Pseudo R-squ.:** | 0.1865 | **Log-Likelihood:** | -37.285 | | | |
| **converged:** | True | **LL-Null:** | -45.835 | | | |
| **Covariance Type:** | nonrobust | **LLR p-value:** | 0.0006742 | | | |
| **Features** | **coef** | **std err** | **z** | **P>\|z\|** | **[0.025** | **0.975]** |
| **Avg Fad Dist** | -0.6903 | 0.514 | -1.344 | 0.179 | -1.697 | 0.317 |
| **Avg Fad Angle** | 0.0682 | 0.043 | 1.582 | 0.114 | -0.016 | 0.153 |
| **Avg Docking Energy Efficiency** | -1.4247 | 0.417 | -3.417 | 0.001 | -2.242 | -0.608 |
| **Avg Pafnucy pKd Efficiency** | -12.1392 | 5.697 | -2.131 | 0.033 | -23.306 | -0.972 |

Table S4: Predicted reactivity classification report against known reactivities for consensus logistic regression model and ligand-specific logistic regression models.

| **Average** | **0** | **1** | **accuracy** | **macro avg** |
|---|---|---|---|---|
| **precision** | 0.80 | 0.79 | 0.79 | 0.79 |
| **recall** | 0.69 | 0.87 | 0.79 | 0.78 |
| **f1-score** | 0.74 | 0.83 | 0.79 | 0.78 |
| **support** | 29.00 | 38.00 | 0.79 | 67.00 |

| **2** | **0.00** | **1.00** | **accuracy** | **macro avg** |
|---|---|---|---|---|
| **precision** | 0.76 | 0.74 | 0.75 | 0.75 |
| **recall** | 0.63 | 0.84 | 0.75 | 0.74 |
| **f1-score** | 0.69 | 0.78 | 0.75 | 0.74 |
| **support** | 30.00 | 37.00 | 0.75 | 67.00 |

| **3** | **0** | **1** | **accuracy** | **macro avg** |
|---|---|---|---|---|
| **precision** | 0.77 | 0.75 | 0.76 | 0.76 |
| **recall** | 0.77 | 0.75 | 0.76 | 0.76 |
| **f1-score** | 0.77 | 0.75 | 0.76 | 0.76 |
| **support** | 35.00 | 32.00 | 0.76 | 67.00 |

| **4** | **0** | **1** | **accuracy** | **macro avg** |
|---|---|---|---|---|
| **precision** | 0.74 | 0.80 | 0.76 | 0.77 |
| **recall** | 0.90 | 0.57 | 0.76 | 0.73 |
| **f1-score** | 0.81 | 0.67 | 0.76 | 0.74 |
| **support** | 39.00 | 28.00 | 0.76 | 67.00 |

| **5** | **0** | **1** | **accuracy** | **macro avg** |
|---|---|---|---|---|
| **precision** | 0.75 | 0.70 | 0.75 | 0.73 |
| **recall** | 0.93 | 0.33 | 0.75 | 0.63 |
| **f1-score** | 0.83 | 0.45 | 0.75 | 0.64 |
| **support** | 46.00 | 21.00 | 0.75 | 67.00 |

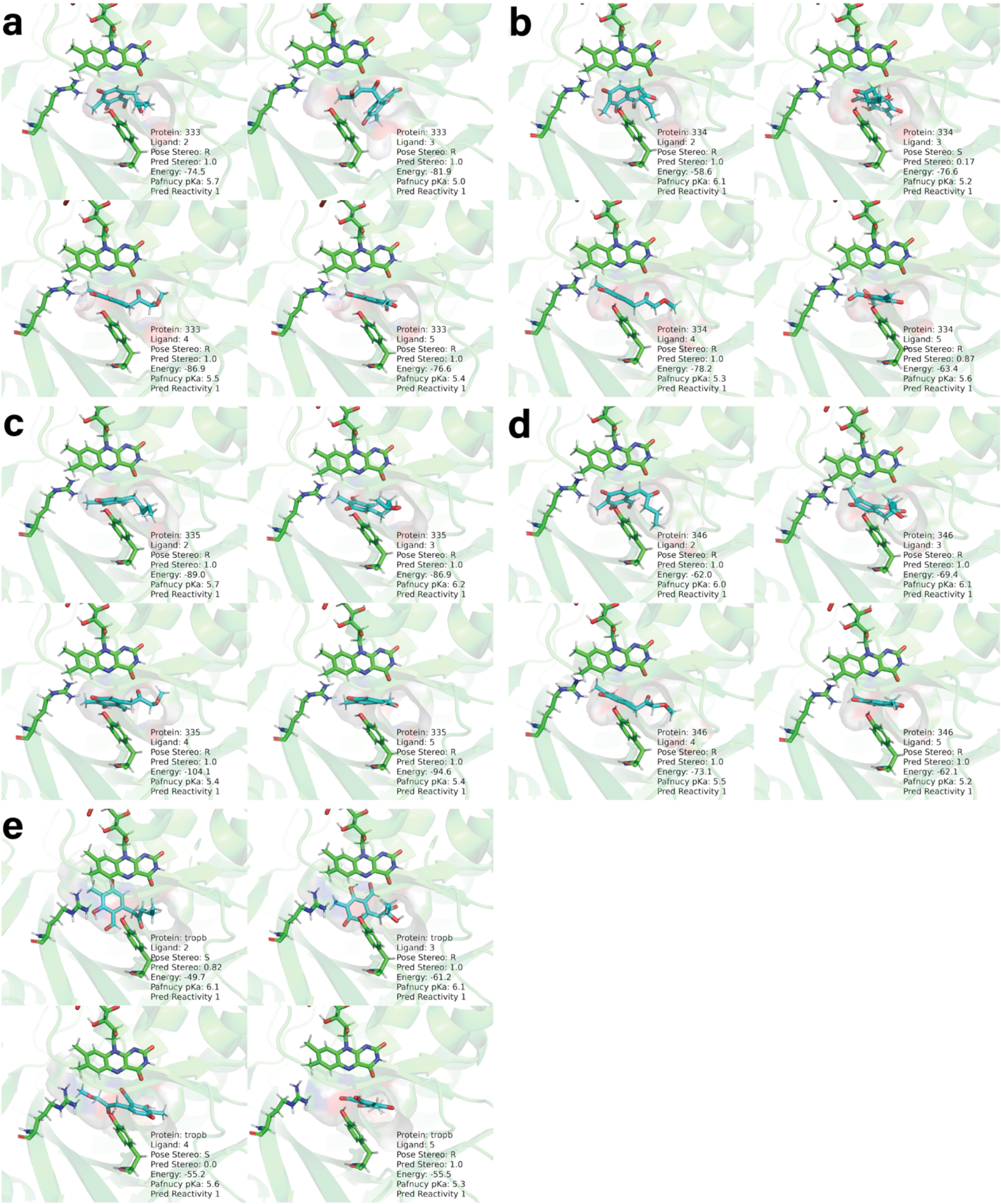

Figure S11: TropB ancestors with R stereochemistry in ancestral FDMO library docked with ligands **2-5**. a)Ancestor 333. b)Ancestor 334. c)Ancestor 335. d)Ancestor 346. e)TropB

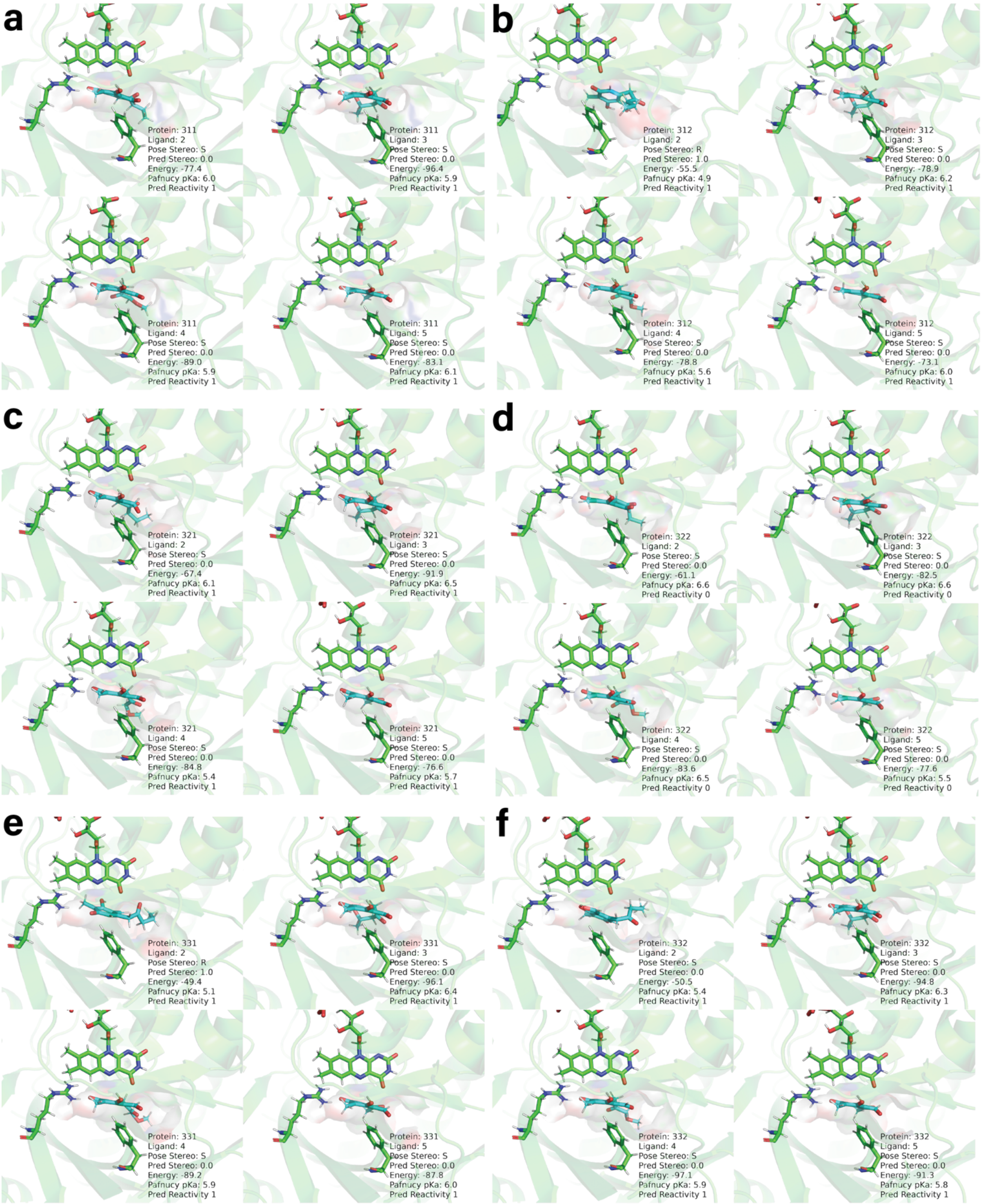

Figure S12: TropB ancestors with S stereochemistry in ancestral FDMO library docked with ligands **2-5**. a) Ancestor 311 (R/S). b) Ancestor 312. c) Ancestor 321. d) Ancestor 322. e) Ancestor 331. f) Ancestor 332.

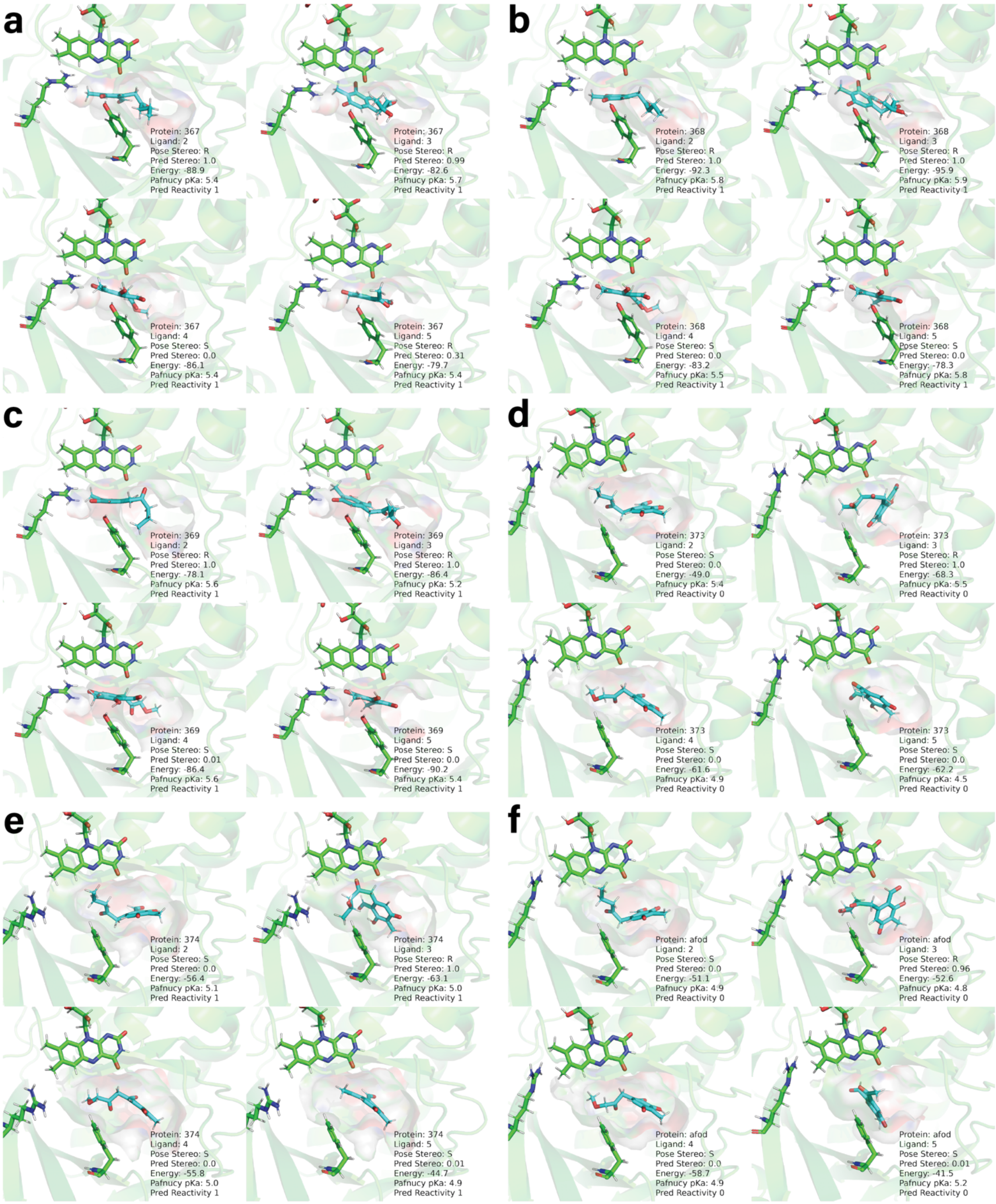

Figure S13: AfoD ancestors with experimentally determined stereochemistries. a) Ancestor 367 (R). b) Ancestor 368 (R). c) Ancestor 369 (R). d) Ancestor 373 (S). e) Ancestor 374 (S). f) AfoD (S).

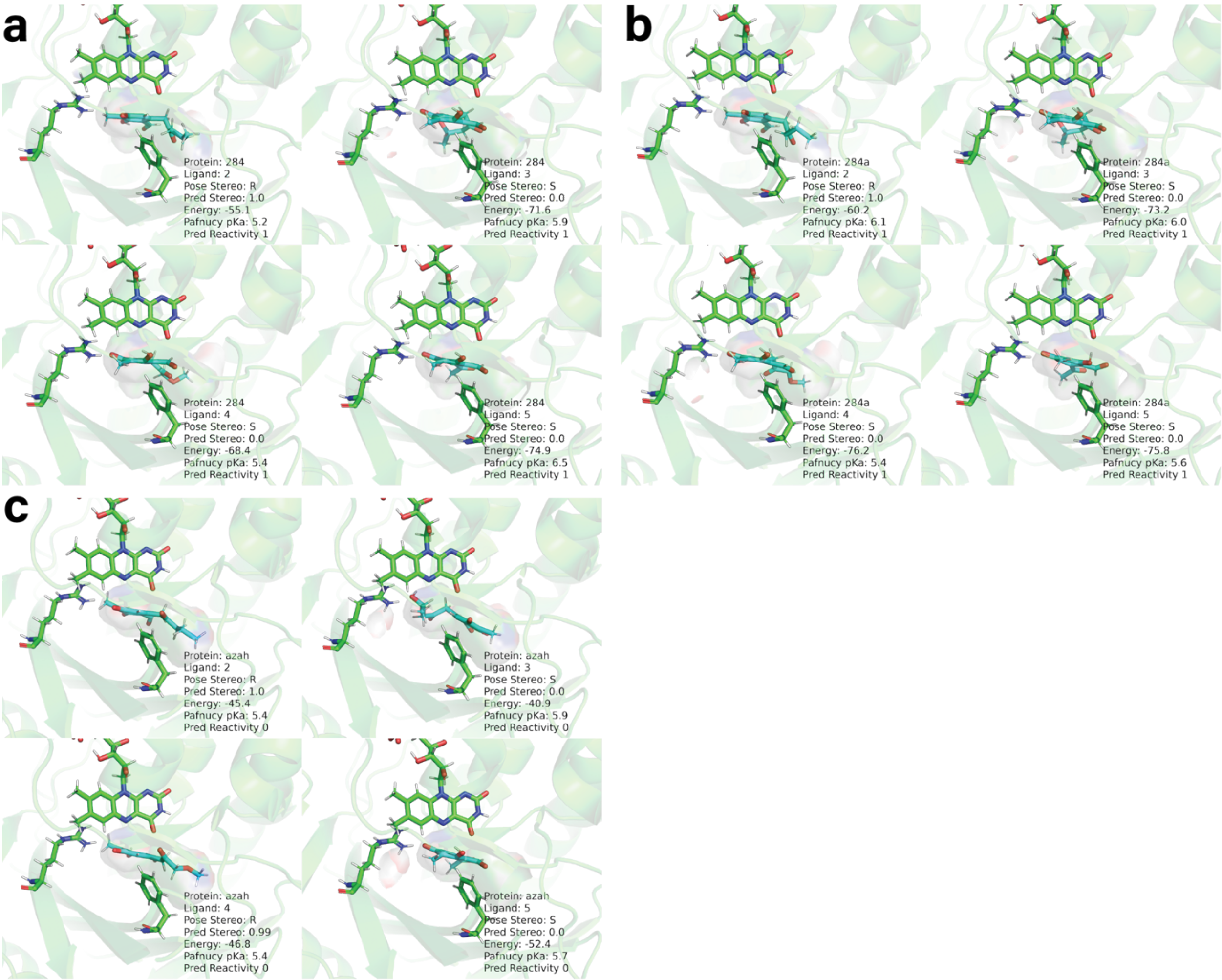

Figure S14: AzaH ancestors with experimentally determined stereochemistries. a) Ancestor 284 (R). b) Ancestor 284a (R). c) AzaH (R).

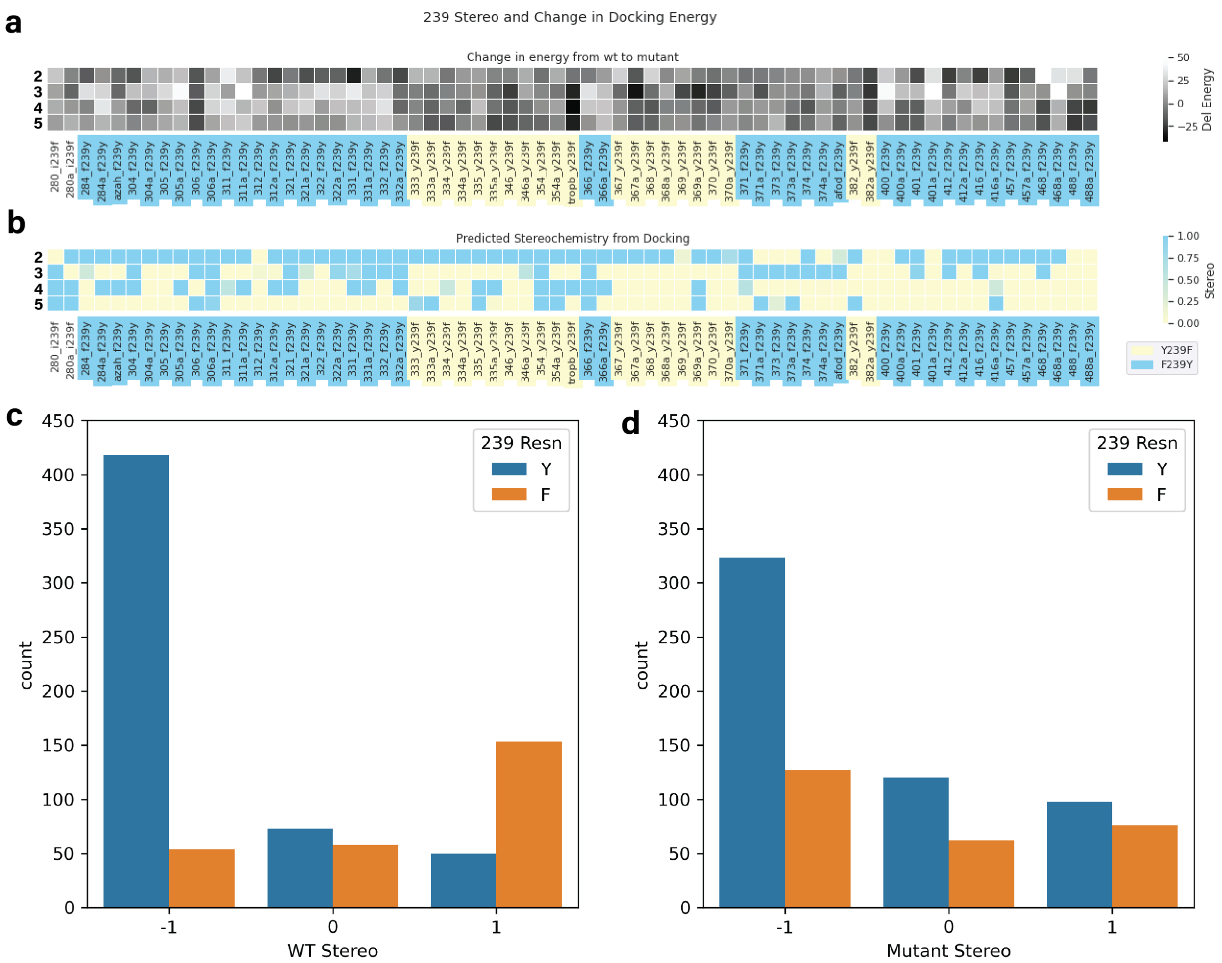

Figure S15: Effect of residue 239 mutations to ancestral FDMO library and full sequence library. a) Docking energy change between mutation and wildtype top docking pose for ancestral FDMO library. F239Y mutations are denoted in blue as they should promote R stereochemistry, and Y239F mutations are denoted in yellow as they should promote S stereochemistry. b) Predicted stereochemistry for ancestral FDMO library, blue squares denote R and yellow squares denote S, with green denoting R/S. c) Wildtype stereochemistries of full sequence library, with blue representing wildtype sequences before F239Y mutation, and orange representing wildtype sequences before Y239F mutation. d) Mutant stereochemistries of full sequence library.

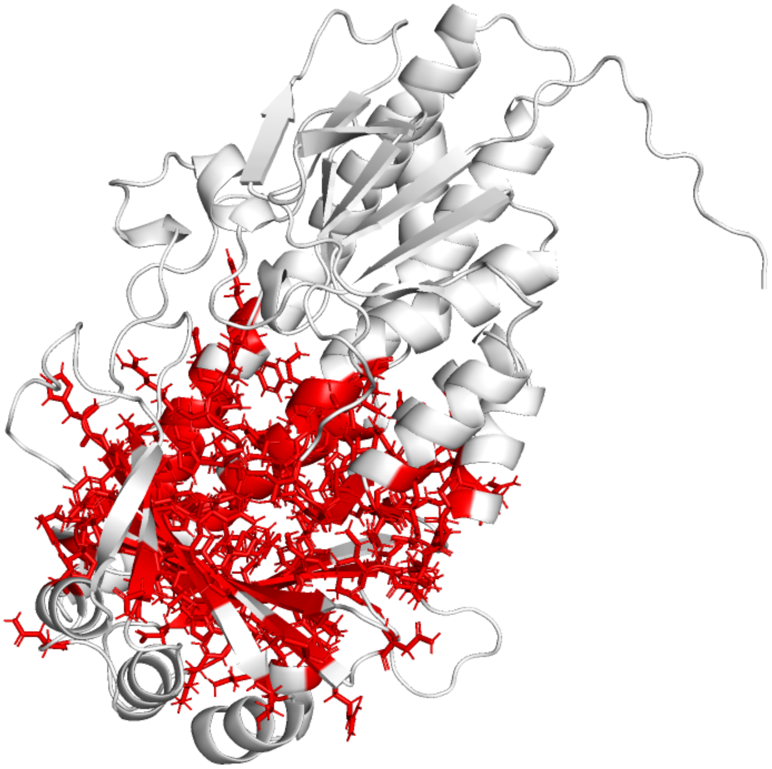

Figure S16: Binding site and second shell residues included after dropping columns with more than 10% gaps represented in red in TropB, excluded residues in white.

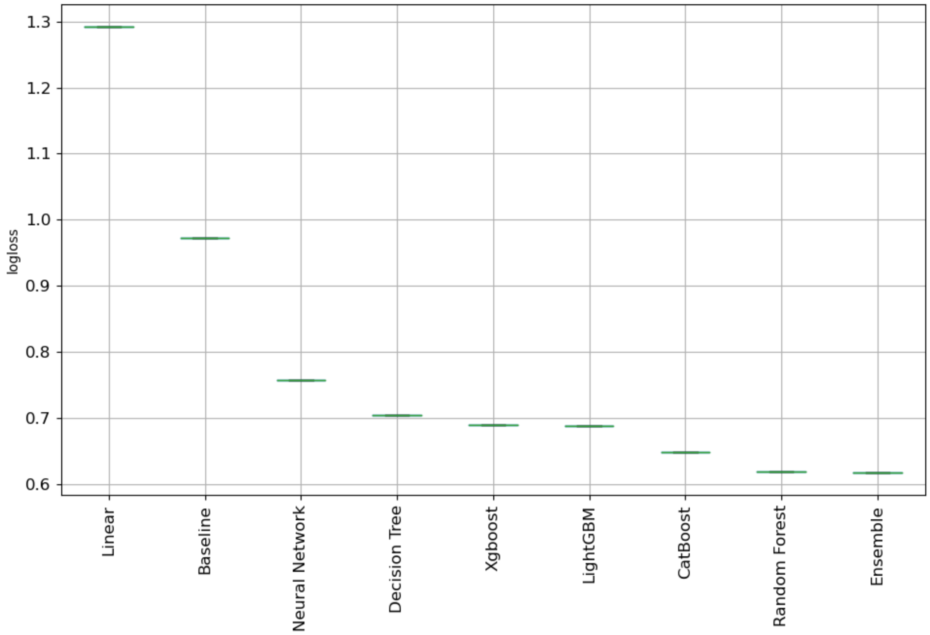

Figure S17: Boxplot of default model performances from MLJAR[18] for stereochemistry prediction on wildtype sequences.

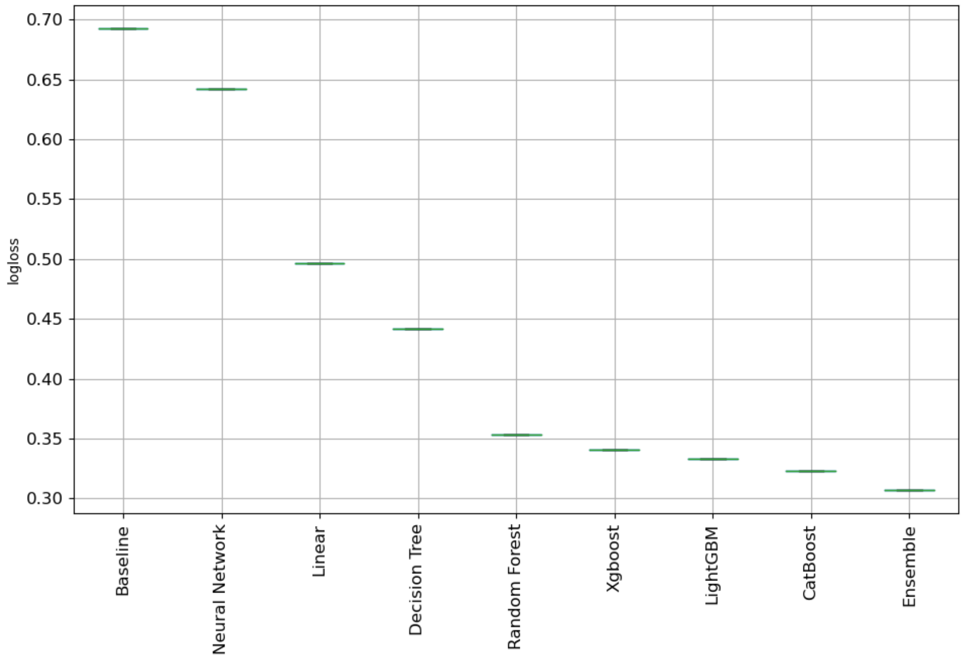

Figure S18: Boxplot of default model performances from MLJAR[18] for conversion prediction on wildtype sequences.

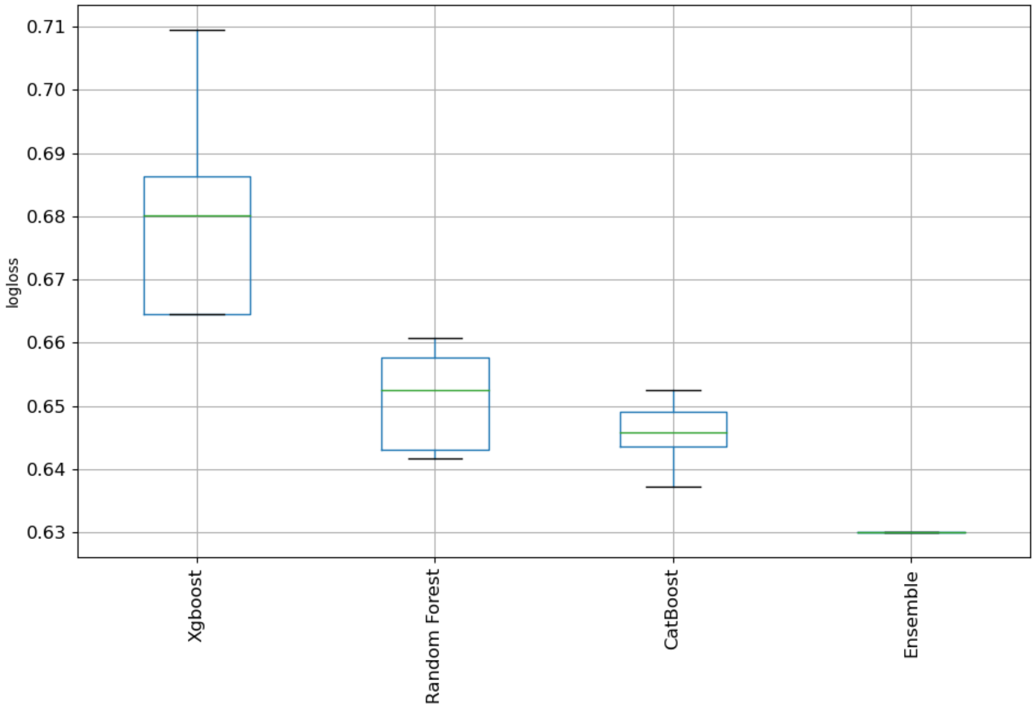

Figure S19: Boxplot of tuned model performances from MLJAR[18] for stereochemistry prediction on wildtype library.

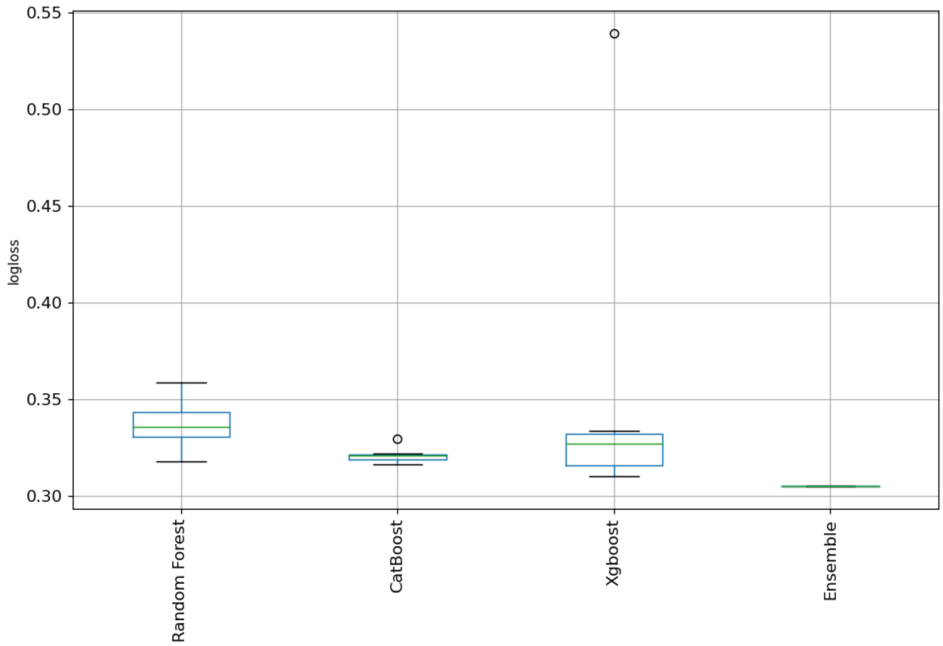

Figure S20: Boxplot of tuned model performances from MLJAR[18] for reactivity prediction on wildtype library.

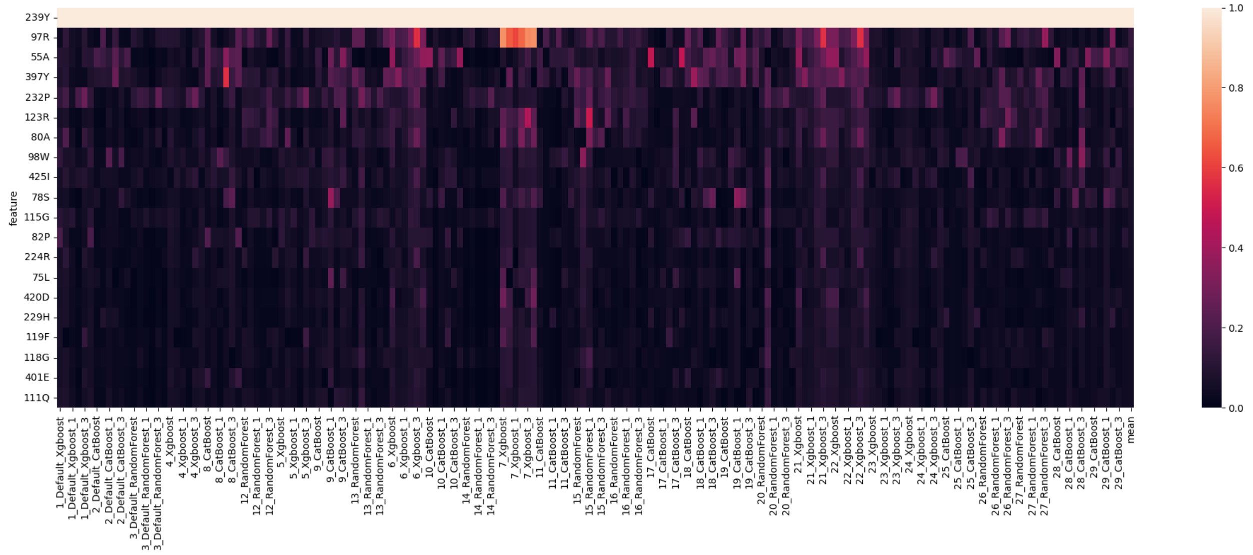

Figure S21: Heatmap of mean SHAP[19] values by residue for every fold of every model for stereochemistry prediction of wildtype sequences.

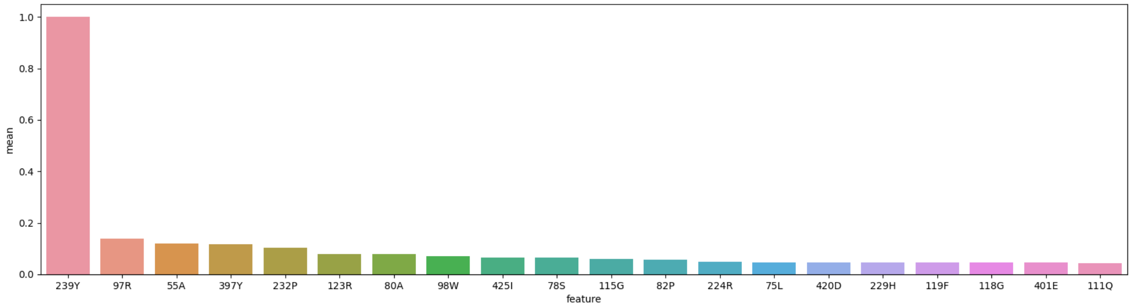

Figure S22: Bar plot of mean SHAP[19] values averaged across all folds/models for stereochemistry prediction of wildtype sequences.

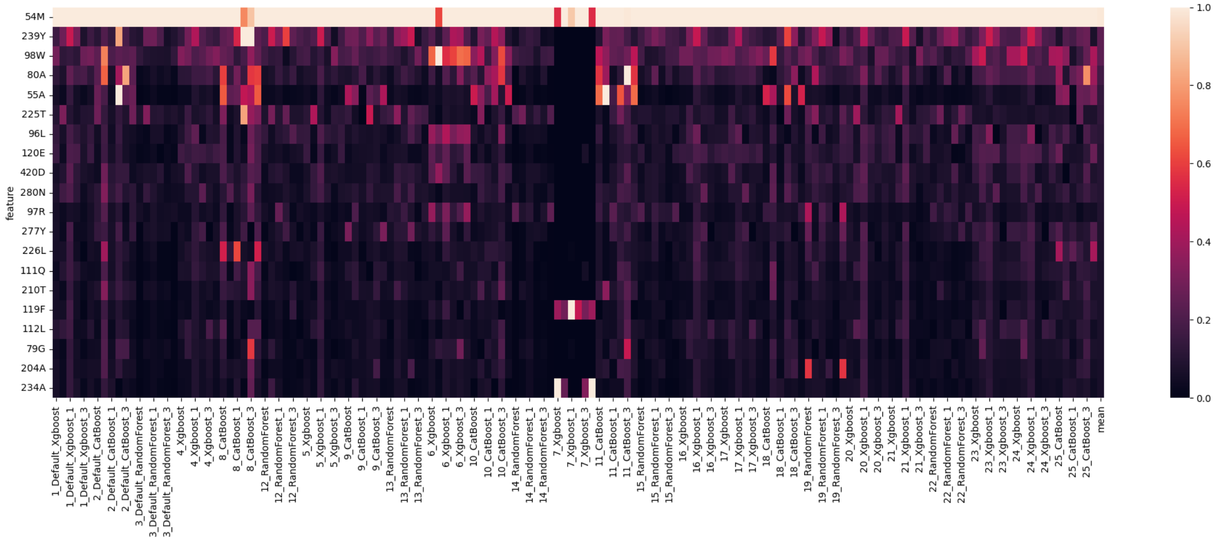

Figure S23: Heatmap of mean SHAP[19] values by residues for every fold of every model for reactivity prediction of wildtype sequences.

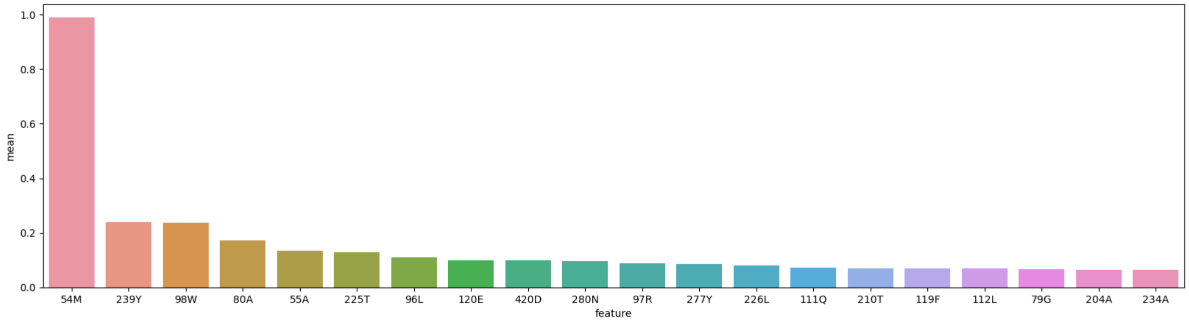

Figure S24: Bar plot of mean SHAP[19] values averaged across all folds/models for reactivity prediction of wildtype sequences

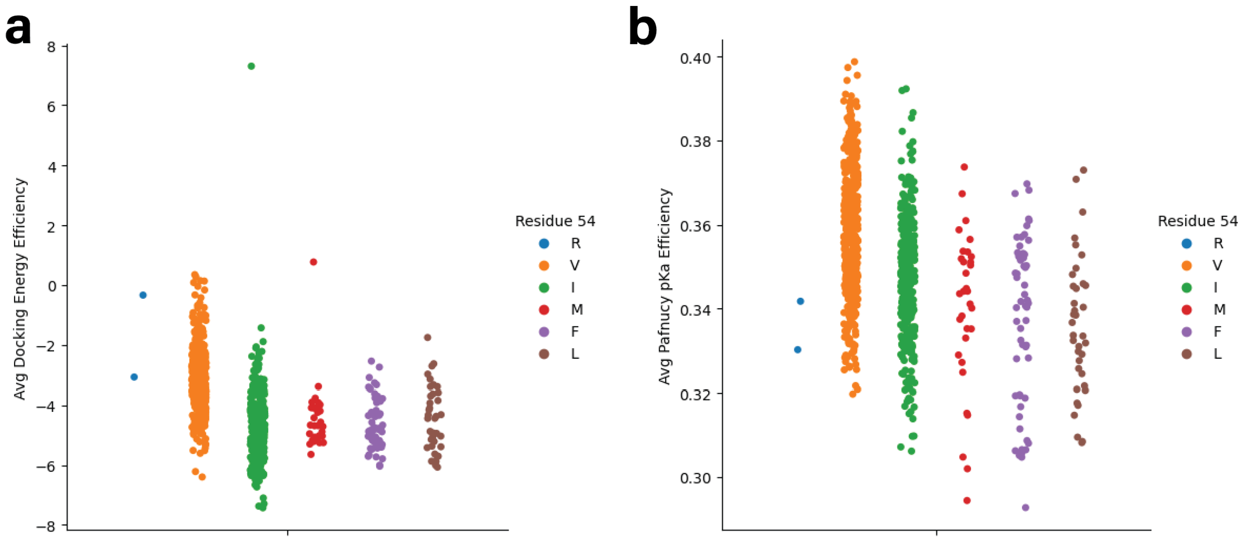

Figure S25: Reactivity metrics and correlation with residue 54. a) Strip plot of average docking energy efficiency across **2-5** separated by amino acid type at residue 54. b) Strip plot of average Pafnucy[17] $pK_d$ efficiency across **2-5** separated by amino acid type at residue 54.

### References

1. Zhang, Y. & Skolnick, J. TM-align: a protein structure alignment algorithm based on the TM-score. doi:10.1093/nar/gki524.
2. Rodríguez Benítez, A. *et al.* Structural Basis for Selectivity in Flavin-Dependent Monooxygenase-Catalyzed Oxidative Dearomatization. *ACS Catal* **9**, 3633–3640 (2019).
3. Feig, M., Karanicolas, J. & Brooks C. L. III. MMTSB Tool Set: enhanced sampling and multiscale modeling methods for applications in structural biology. *J Mol Graph Model* **22**, 377–395 (2004).
4. Brooks, B. R. *et al.* CHARMM: The Biomolecular Simulation Program. *J Comput Chem* **30**, 1545 (2009).
5. Huang, J. *et al.* CHARMM36m: an improved force field for folded and intrinsically disordered proteins. *Nat Methods* **14**, 71–73 (2017).
6. Buckner, J. *et al.* pyCHARMM: Embedding CHARMM Functionality in a Python Framework. *J Chem Theory Comput* **19**, 3752–3762 (2023).
7. Ding, X., Wu, Y., Wang, Y., Vilseck, J. Z. & Brooks C. L. III. Accelerated CDOCKER with GPUs, Parallel Simulated Annealing, and Fast Fourier Transforms. *J Chem Theory Comput* **16**, 3910–3919 (2020).
8. Vanommeslaeghe, K. *et al.* CHARMM General Force Field (CGenFF): A force field for drug-like molecules compatible with the CHARMM all-atom additive biological force fields. *J Comput Chem* **31**, 671 (2010).
9. Chiang, C.-H. *et al.* Deciphering the evolution of flavin-dependent monooxygenase stereoselectivity using ancestral sequence reconstruction. *Proceedings of the National Academy of Sciences* **120**, e2218248120 (2023).
10. Jumper, J. *et al.* Highly accurate protein structure prediction with AlphaFold. *Nature 2021 596:7873* **596**, 583–589 (2021).

11. Yang, J. & Zhang, Y. I-TASSER server: new development for protein structure and function predictions. *Web Server issue Published online* **43**, (2015).
12. Källberg, M. *et al.* Template-based protein structure modeling using the RaptorX web server. *Nat Protoc* **7**, 1511–1522 (2012).
13. Y, S. *et al.* High-resolution comparative modeling with RosettaCM. *Structure* **21**, 1735–1742 (2013).
14. Baek, M. *et al.* Accurate prediction of protein structures and interactions using a three-track neural network. *Science (1979)* **373**, 871–876 (2021).
15. Waterhouse, A. *et al.* SWISS-MODEL: homology modelling of protein structures and complexes. *Nucleic Acids Res* **46**, W296–W303 (2018).
16. Mariani, V., Biasini, M., Barbato, A. & Schwede, T. lDDT: a local superposition-free score for comparing protein structures and models using distance difference tests. *Bioinformatics* **29**, 2722–2728 (2013).
17. Stepniewska-Dziubinska, M. M., Zielenkiewicz, P. & Siedlecki, P. Development and evaluation of a deep learning model for protein-ligand binding affinity prediction. *Bioinformatics* **34**, 3666–3674 (2018).
18. Plonska, A. & Plonski, P. MLJAR: State-of-the-art Automated Machine Learning Framework for Tabular Data.  . Preprint at https://github.com/mljar/mljar-supervised (2021).
19. Lundberg, S. M., Allen, P. G. & Lee, S.-I. A Unified Approach to Interpreting Model Predictions. in *Advances in Neural Information Processing Systems 30* 4765–4774 (2017).
